# Ketones uncouple Alzheimer’s disease pathology and cognition from metabolic health in two mouse models

**DOI:** 10.64898/2026.11.24.768721

**Authors:** Michelle M. Sonsalla, Reji Babygirija, Sunny Gupta, Alexandra Trebil, Brenna Virnig, Nathan Buckley, Bailey A. Knopf, Mari Cole, Madeline Johnson, Isaac Grunow, Fan Xiao, Yang Liu, Diana Vertein, Michael J. Rigby, Chung-Yang Yeh, Snehal N. Chaudhari, David A. Harris, Luigi Puglielli, Dudley W. Lamming

## Abstract

Ketogenic diet (KD), an extremely low carbohydrate diet that induces ketone production, has shown some promise as a treatment for Alzheimer’s disease (AD) in both human studies and mouse models. Here, we examined the effects of a cyclic KD (cKD) on metabolic health in the APP/PS1 and 3xTg mouse models of AD. While improvements in metabolic health are coupled with reduced AD pathology and improved cognition in many other AD interventions, we find that cKD improves AD pathology and cognition in a sex- and strain-specific manner despite somewhat negative consequences for metabolic health. Boosting ketone production with the dietary supplement 1,3-butanediol similarly impaired metabolic health while improving cognition. Finally, we found that ketone-induced benefits to cognition and AD pathology may be partly construed by alterations to mTOR signaling and autophagy. In summary, while the specific benefits of increased ketones vary between mouse models, sexes, and the specific diet or supplement used, increased ketones can slow or prevent AD pathology and preserve cognition without necessarily improving metabolic health. This work has important implications for ketone-based therapies, as the individual response to KD may vary, and ketone supplements may not fully recapitulate the benefits of KD on AD development and progression.

## Introduction

An estimated 7.4 million individuals in the United States are living with Alzheimer’s disease (AD) (2026). Of these, almost two-thirds are women, and many studies have shown differences in the pathology of AD between men and women (Filon, Intorcia et al. 2016, Aggarwal and Mielke 2023, Castro-Aldrete, Moser et al. 2023), highlighting the importance in studying the impact of possible interventions in both sexes. While some FDA-approved pharmaceutical interventions for AD exist (Fox, Belder et al. 2025), the options are limited in scope, expensive, and associated with many possible adverse side effects (Ross, Weinberg et al. 2022).

Dietary interventions, which are more accessible, are an attractive alternative to costly pharmaceutical interventions. AD is associated with negative metabolic health in both humans and rodents (Barron, Rosario et al. 2013, Knight, Martins et al. 2014, Vandal, White et al. 2015, Velazquez, Tran et al. 2017, Robison, Gannon et al. 2020, Sullivan 2020, Wieckowska-Gacek, Mietelska-Porowska et al. 2021, Mietelska-Porowska, Domanska et al. 2022, Abubakar, Giri et al. 2025), and many interventions that improve metabolic health also show improvements to AD pathology and cognition (Patel, Gordon et al. 2005, Wang, Ho et al. 2005, Halagappa, Guo et al. 2007, Parrella, Maxim et al. 2013, Schafer, Alldred et al. 2015, Elias, Padinjakara et al. 2023, Babygirija, Sonsalla et al. 2024, Sonsalla, Babygirija et al. 2025, M’Bra, Hamilton et al. 2026). There is growing interest in using geroprotective interventions for the treatment of AD, and a number of dietary interventions that promote healthy aging in wild-type mice (Newman, Covarrubias et al. 2017, Pak, Haws et al. 2021, Hill, Albarado et al. 2022, Green, Trautman et al. 2023, Di Francesco, Deighan et al. 2024) have been shown to slow or prevent AD in mouse models of the disease (Halagappa, Guo et al. 2007, Babygirija, Sonsalla et al. 2024, Babygirija, Han et al. 2025, Babygirija, Green et al. 2026).

Ketogenic diet (KD) was originally established as a treatment for some types of drug-resistant epilepsy (Wilder 1921, Peterman 1925). When blood glucose levels are low, ketogenesis generates ketones as a vital, readily utilizable energy source for many organs in the body, including the brain. Many studies have shown that KD can improve brain metabolism, AD pathology, and cognition (Van der Auwera, Wera et al. 2005, Newman, Covarrubias et al. 2017, Roberts, Wallace et al. 2017, Xu, Jiang et al. 2022, Di Lucente, Persico et al. 2024, Abdel-Aziz, Ahmed et al. 2025, Cimen, Elibol et al. 2025, Di Lucente, Rutkowsky et al. 2026). As glucose utilization in the brain is often impaired in AD (Friedland, Budinger et al. 1983, Azari, Pettigrew et al. 1993, Minoshima, Giordani et al. 1997, Mosconi 2005, Correia, Santos et al. 2011, Chen and Zhong 2013), it has been suggested that a KD might help maintain proper brain function by providing an alternative fuel source. In addition, ketones can act as HDAC inhibitors, HCA2 agonists, and modulators of neurotransmitters, providing multiple possible mechanisms for a KD to improve AD pathology and cognition (reviewed in (Newman and Verdin 2014, Newman and Verdin 2017)).

Furthermore, both KD and ketones themselves inhibit the nutrient sensing protein kinase mechanistic target of rapamycin complex 1 (mTORC1)(McDaniel, Rensing et al. 2011, Willemse, Terburgh et al. 2023, Matawali, Yeap et al. 2025, Wang, Hou et al. 2026), providing another possible mechanism for improvements in AD pathology. mTORC1 hyperactivation is a common attribute of age-related diseases, including AD (Caccamo, Majumder et al. 2010, Gouras 2013, O’ Neill 2013, Cai, Chen et al. 2015, Perl 2015, Romero, Bueno et al. 2016, Gonzales 2019, Bloom and Norambuena 2023, Cai, Hou et al. 2023), and can inhibit autophagy, leading to reduced clearance of amyloid and subsequent aggregation into plaques (Nixon, Wegiel et al. 2005, Pickford, Masliah et al. 2008, Salminen, Kaarniranta et al. 2012, Salminen, Kaarniranta et al. 2013). Prior work has shown that mTORC1 inhibition via rapamycin or dietary intervention has been sufficient to increase autophagy and improve AD pathology and cognition (Caccamo, Majumder et al. 2010, Spilman, Podlutskaya et al. 2010, Majumder, Richardson et al. 2011, Babygirija, Sonsalla et al. 2024, Babygirija, Green et al. 2025), suggesting that KD may also improve AD pathology and cognition via modulation of mTORC1 and autophagy.

However, there are hurdles to implementing KD as a translatable intervention, as KD may be associated with hyperlipidemic side effects and has low long-term adherence (Buren, Ericsson et al. 2021, Gallop, Vieira et al. 2025). Additionally, KD in mice drastically impairs metabolic health (Gallop, Vieira et al. 2025). Therefore, it is important to examine alternative treatment options that may convey the benefits of KD without negative metabolic consequences or the need for dietary changes. Recently, a cyclic KD (cKD) paradigm has been utilized to avoid the negative metabolic effects of KD while improving lifespan and age-related memory decline (Newman, Covarrubias et al. 2017). Exogenous ketone supplementation are another potential option. Two promising supplements are a β-HB precursor, 1,3-butanediol (1,3-BD) and medium chain triglycerides (MCT). Prior work has shown potential for these supplements in AD mouse models, including improvements in cognition and AD pathology (Chen, Li et al. 2025, Fulghum, Hayir et al. 2026, M’Bra, Hamilton et al. 2026).

In the present study, we utilize both females and males of two different AD mouse models, APP/PS1 and 3xTg, to examine the role of both sex and strain on the response to a KD. APP/PS1 mice express a chimeric mouse/human APP_swe_ and a mutant human PSEN1_dE9_, resulting in robust amyloid pathology and age-related worsening of amyloid-β (Aβ) load accompanied by synapse loss, gliosis, and cognitive impairments (Jankowsky, Fadale et al. 2004, Kamphuis, Mamber et al. 2012, Hong, Beja-Glasser et al. 2016). 3xTg mice express human APP_swe_, tau_P301L_, and PSEN1_M146V_, and thus develop both amyloid and tau pathology (Oddo, Caccamo et al. 2003, Oddo, Caccamo et al. 2003); these mice show age-related increases in Aβ levels, tau phosphorylation, synapse loss, gliosis, and cognitive deficits (Oddo, Caccamo et al. 2003, Oddo, Caccamo et al. 2003, Belfiore, Rodin et al. 2019). Using both models allows us to more comprehensively assess the effect of KD on AD pathology and cognition.

Here, we placed APP/PS1 and 3xTg mice onto either a control or cKD paradigm and assessed metabolic health, cognition, and AD pathology. Overall, we found that cKD improved body composition in 3xTg mice but worsened glucose homeostasis in both strains. KD improved cognition in both APP/PS1 and 3xTg mice, with the degree and type of improvement differing by sex and strain. Amyloid pathology was improved in APP/PS1 mice, while 3xTg females showed reduced Tau phosphorylation. KD reduced mTORC1 signaling in both strains, but we observed strain-specific effects on autophagy. We find that supplementing 3xTg mice with 1,3-BD has beneficial effects for AD; however, neither supplement recapitulated all of the benefits of a KD. These results show that dietary ketones are a promising intervention for AD, but the benefits of a KD for the development and progression of AD are highly sex- and strain-dependent and not fully recapitulated by ketone supplements. This work has important implications for administering ketone-based therapies, as the response to KD may vary between individuals, and ketone supplements may not fully recapitulate the benefits of KD on AD development and progression.

## Materials and Methods

### Animals

All procedures were performed in accordance with institutional guidelines and were approved by the Institutional Animal Care and Use Committee (IACUC) of the William S. Middleton Memorial Veterans Hospital (Madison, WI, USA) and the University of Wisconsin-Madison IACUC. All mice were maintained at a temperature of approximately 22°C, with a 12:12 light/dark cycle with food and water available ad libitum. Health checks were completed on all mice daily. Mice were acclimatized to our experimental facility for one week before experiment start and were housed 2-3 per cage.

Mice for all experiments were originally obtained from The Jackson Laboratory/Mutant Mouse Resource and Research Centers (MMRC). Homozygous male and female 3xTg mice (Stock #034830) were bred together to produce mice used for experiments. Hemizygous male APP/PS1 mice (Stock #034829) were bred against non-transgenic controls to maintain a colony containing hemizygous APP/PS1 mice and non-transgenic littermates. Diets for all experiments were obtained from Inotiv (formerly Envigo), and full descriptions of each diet are provided in **Table S1**. Randomization of mice to groups was performed at cage level to ensure all groups had approximately the same starting weight and body composition.

For the first experiment, 4-month-old male and female APP/PS1 mice were randomly assigned to one of two diet conditions: Control (Con) or cyclic ketogenic diet (cKD) (diet compositions in **Table S1**). Briefly, the Con diet (TD.220339; same composition as TD.150345) consists of approximately 10% protein, 78% carbohydrates, and 12% fat while KD (TD.160153) is made up of approximately 10% protein and 90% fat. These diets have been previously used in numerous studies (Newman, Covarrubias et al. 2017, Tognini, Murakami et al. 2017, Ang, Alexander et al. 2020, Nomura, Murad et al. 2024). The cKD paradigm consisted of 1-week cycles between the Control and Ketogenic diets. Mice were maintained on these diets for 24 weeks. In the second experiment, 6-month-old male and female 3xTg mice were maintained on either control or cKD diets for 28 weeks to assess the impact of cKD on an AD model containing more robust tau pathology. Finally, in the third experiment 6-month-old male and female 3xTg mice were maintained on either control diet or diets containing 1,3-butanediol (TD.160257; 10% 1,3-Butanediol; Sigma Aldrich, St. Louis, MO, USA) (Tognini, Murakami et al. 2017) or medium chain triglycerides (TD.230552; 2% caprylic acid; Sigma Aldrich, St. Louis, MO, USA) (comparable to those used in previous studies (Zhang, Xue et al. 2019, Zhang, Zhang et al. 2022, Kawahara, Fujiwara-Tani et al. 2025)) for 28 weeks to assess the ability of these compounds to increase ketone levels and effect AD pathology independent of KD.

### In vivo procedures

Food consumption in home cages was measured by moving mice to clean cages, filling the hopper with a measured quantity of fresh diet, and measuring the remainder 3-4 days later. The amount of food consumed was adjusted for the number of mice in each cage, the number of days that had passed, and the relative weights of the mice.

Mouse body composition was analyzed at several points throughout the study using an EchoMRI Body Composition Analyzer (EchoMRI, Houston, TX, USA). Metabolic parameters (O_2_, CO_2_, food composition, respiratory exchange ratio (RER), and energy expenditure) and activity tracking were analyzed using a Columbus Instruments Oxymax/CLAMS-HC metabolic chamber system. Mice were acclimated to the chamber cages for ∼24 hours and data from a subsequent, continuous 24-hour period was recorded and analyzed.

Blood ketone levels were measured using a KetoBM blood ketone meter and test strips (KetoBM, Sheridan, WY, USA). Glucose tolerance tests were performed by fasting the mice overnight for 16 hours then injecting glucose (1 g kg-1) intraperitoneally (I.P.) (Yu, Yang et al. 2018, Bellantuono, de Cabo et al. 2020). Insulin tolerance tests were performed by fasting the mice for 4 hours then injecting insulin I.P. (0.75 U kg-1). Blood glucose measurements were taken using a Bayer Contour Next blood glucose meter (Bayer, Leverkusen, Germany) and test strips.

Behavioral assays were performed on all mice and recorded/analyzed using the Ethovision XT Animal Tracking Software by Noldus. Before each procedure mice were allowed to acclimate to the behavioral testing room for 30 minutes. Between each run the arenas and objects were cleaned with 70% ethanol to remove olfactory cues.

For dark/light box, mice were placed into the dark portion of a box that was 1/3 enclosed and 2/3 open, with a small opening for movement between the two zones. The mice were allowed to explore the box for 5 minutes, during which time latency to entry into light zone and cumulative time spent in light zone were measured.

For Novel Object Recognition (NOR), mice were first habituated to the empty arena for 5 minutes with no objects, which also acted as the open field test. Approximately 24 hours later, an acquisition trial was performed where the mice were placed in the arena with two identical objects (hereafter referred to as “Object A”), equidistant from the mouse. The mouse was allowed to explore the two objects for 5 minutes before returning to the home cage. One hour after the acquisition trial, a short-term memory (STM) test was performed during which the mouse was placed into the arena with one of object A as well as a single novel object, or “Object B”, and allowed to explore for 5 minutes. The following day, 24 hours after the STM test, a long-term memory (LTM) test was performed. During the LTM test, the mouse was placed in the arena with one of object A and a single novel object, “Object C”, and allowed to explore for 5 minutes. Results were quantified by calculating discrimination index (DI) by dividing the time spent exploring the novel object by the time spent examining both objects. A DI value between -0.2 and 0.2 indicates no difference in time spent examining each object. A positive value indicates greater time was spent examining the novel object whereas a negative value indicates greater time was spent examining the familiar object. Due to the nature of the DI calculation, absolute DI is also calculated to separate discrimination from object preference.

Barnes maze testing involved 4 phases: habituation, acquisition, and STM and LTM memory tests. During habituation, the mice were placed on the maze and immediately led to the escape box where they remained for 2 minutes before returning to their home cage. After approximately 15 minutes, the mice underwent their first acquisition trial in which they were allowed to explore the maze until either they found the escape box or 3 minutes had elapsed. If the mouse did not find the escape box in 3 minutes, they were gently led to the escape box. Regardless of how the mouse reached the escape box, they remained there for 1 minute after every trial. Over the first four days, four acquisition trials were performed each day with an inter-trial interval of 15 minutes between each trial. On days 5 (STM test) and 12 (LTM test), the escape box was blocked and the mouse was placed on the maze for 90s to determine how well it was able to recall where the escape box was. Latency to goal was analyzed for all trials and average for each day of training and testing.

Mice were euthanized by cervical dislocation after a 3-hour fast at the beginning of the light cycle and tissues for molecular analysis were either flash-frozen in liquid nitrogen and stored at -80°C, or fixed and prepared as described below. For brain histology, one hemisphere of the brain was collected and fixed in 15% neutral buffered formalin for 24 hours before transferring to 70% ethanol for storage until later analysis. In the cKD cohort, half of the mice were euthanized at the end of the KD^off^ period and half at the end of the KD^on^ period.

### Histology

Formalin-fixed half brain samples were paraffin-embedded by the University of Wisconsin Carbone Cancer Center Experimental Animal Pathology Laboratory (UWCCC EAPL). 5µm sections were then cut using a Leica microtome (Wetzlar, Germany) and affixed to Superfrost Plus microscope slides (Thermo Fisher Scientific, Waltham, MA, USA). For amyloid plaque staining, brain sections were deparaffinized and rehydrated, then treated in 70% formic acid for epitope retrieval. Sections were blocked with normal goat serum (NGS) and incubated with 6E10 monoclonal antibody (1:100; #803004; Biolegend, San Diego, CA, USA) overnight before incubation with biotinylated secondary antibody and visualization with diaminobenzidine chromagen. Glial staining was performed by deparaffinizing and rehydrating sections followed by antigen retrieval using a 1x Tris-EDTA (pH9.0) antigen retrieval buffer (Abcam, Cambridge, UK) and stained (1:1000) with anti-Iba1 (ab178847; Abcam, Cambridge, UK) and anti-GFAP (PIMA512023; Thermo Fisher Scientific, Waltham, MA, USA) antibodies. Co-staining of amyloid plaques, microglia, and astrocytes was performed using anti-Iba1 and anti-GFAP as above while also incubating with MethoxyX04 and ToPro3 during secondary antibody staining to stain amyloid plaques and nuclei, respectively. All sections were imaged using an EVOS FL Auto microscope (Thermo Fisher Scientific, Waltham, MA, USA) at 20x magnification. ImageJ was used for quantification by converting images to binary images via an intensity threshold and quantifying positive area.

### Immunoblotting

Brain hemisphere samples were lysed in cold RIPA buffer supplemented with phosphatase inhibitor and protease inhibitor cocktail tablets (Thermo Fisher Scientific, Waltham, MA, USA) using a FastPrep 24 (M.P. Biomedicals, Santa Ana, CA, USA) with bead-beating tubes (VWR, Radnor, PA, USA) and zirconium ceramic oxide beads (Thermo Fisher Scientific, Waltham, MA, USA). Protein lysates were then centrifuged at 13,300 rpm for 10 min and the supernatant was collected. Protein concentration was determined by Bradford Assay (Pierce Biotechnology, Waltham, MA, USA). 20-60 μg protein was separated on 16% tris-glycine gels (ThermoFisher Scientific, Waltham, MA, USA) and transferred to PVDF membrane (EMD Millipore, Burlington, MA, USA). mTORC1 signaling was assessed by downstream targets T389 pS6K and S240/244 pS6. Autophagy markers were assessed, including autophagy proteins Atg5 and Atg7, autophagosome formation protein Beclin, and the autophagy receptor (sequestosome 1, SQSTM1). Tau phosphorylation was assessed using the phospho-T231 Tau antibody. Antibody vendors, catalog numbers and the dilution used are provided in **Table S2**. Imaging was performed using a Bio-Rad Chemidoc MP imaging station (Bio-Rad, Hercules, CA, USA). Quantification was performed by densitometry using NIH ImageJ software.

### ELISA

Amyloid-β 40 (Aβ40) and 42 (Aβ42) in the brain was quantified using human Aβ40 and Aβ42 ELISA kits (Invitrogen, USA, #KHB3482 and #KHB3441, respectively) in accordance with manufacturer’s instructions.

### Metabolite extraction and LC-MS Analysis

Extraction and quantification of metabolites from brain tissue were performed using a previously established protocol (Williams, Kim et al. 2026). Frozen whole brain samples were homogenized in extraction buffer (20 mM ascorbic acid, 20 mM ammonium acetate, and 20 mM β-mercaptoethanol (BME), pH 8.1. 5 μl of pooled rat serum (Santa Cruz Biotechnology, Dallas, TX, USA) was added to tubes containing the supernatant to deconjugate polyglutamylated folates. Supernatants were filtered by SPE using a Strata-X 33-μm polymeric reverse-phase 96-well SPE plate (Phenomenex, Torrance, CA, USA). LC-MS analysis was carried out on a Vanquish ultra-high-performance liquid chromatography system (ThermoFisher Scientific, Waltham, MA, USA) integrated with an Orbitrap Exploris 120 mass spectrometer (ThermoFisher Scientific, Waltham, MA, USA). Metabolite separation was achieved on a reversed-phase polar C18 column (Phenomenex, Torrance, CA, USA) kept at 40 °C using a 12 min gradient of solvent A (95% H2O/5% methanol, 5 mM DMHA, pH 8.1 adjusted with formic acid) and solvent B (methanol containing 5 mM DMHA) at 0.4 mL/min. Eluted compounds were analyzed by heated electrospray ionization mass spectrometry in negative ion mode. Quantification was done by integrating peak areas using MAVEN (version 2.10.21) (Clasquin, Melamud et al. 2012). Signal intensity was normalized to the internal standards (d2-folic acid and methotrexate) and then to tissue weight.

### Statistical Analyses

All statistical analyses were conducted using Prism, version 10 (GraphPad Software Inc., San Diego, CA, USA). Tests involving multiple factors were analyzed by a two-way analysis of variance (ANOVA) with diet and sex as variables, followed by multiple comparisons using Dunnett’s or Sidak’s post-hoc tests as specified in the figure legends. In all figures: asterisks denote differences between diets with * = p<0.05, ** = p<0.01, *** = p<0.001, and **** = p<0.0001.

## Results

### Ketogenic diet alters metabolic health in APP/PS1 mice

Female and male APP/PS1 mice were placed onto either Control (Con; 10% protein, 78% carbohydrates, and 12% fat) or cyclic ketogenic diet (cKD; 1 week cycles of Con diet and KD diet: 10% protein and 90% fat) (diet compositions in **Table S1**) from 4 to 10 months of age, and metabolic health and cognitive performance was assessed (**Fig. 1A**). A cKD was used as cycling on and off of a KD ameliorate metabolic impairments while maintaining KD-related improvements to memory in C57BL/6J mice (Newman, Covarrubias et al. 2017); however, no studies to-date have studied if cKD can slow or prevent AD in mice. We confirmed cKD increased ketone levels, in the refed state while mice were on KD (KD^on^; **Fig. 1B**). Fasted female, but not male, cKD-fed mice also have increased ketone levels while off KD (KD^off^; **Fig. 1B**).

**Figure 1:**
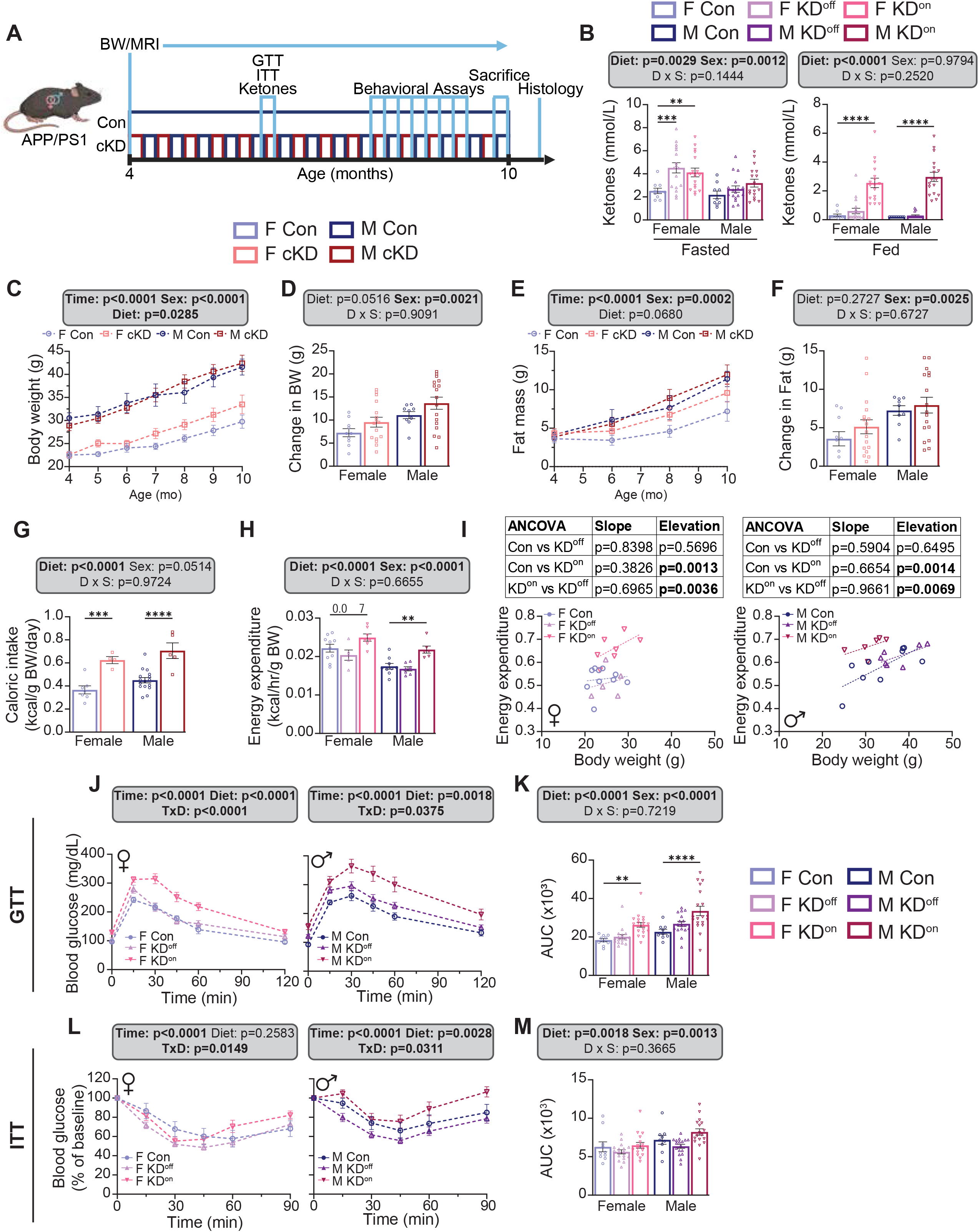
Ketogenic diet worsens glucose homeostasis in APP/PS1 mice. A) Experiment Schematic B) Fasted and fed ketone levels C-D) Body weight across the experimental period (C) and change in body weight from 4 to 10 months (D). E-F) Fat mass across the experimental period (E) and change in fat mass from 4 to 10 months (F). G) Home cage food consumption normalized to body weight. H) Energy expenditure normalized to body weight. I) ANCOVA of energy expenditure. J-K) i.p. glucose tolerance test (J) and area under the curve (K). L-M) i.p. insulin tolerance test (L) and area under the curve (M). (A-F, H-M) Body weight/composition, metabolic chamber, and GTT/ITT data n=9-17/group; (G) home cage data n=4-16/group. Asterisks denote significant differences between diets: *p<0.05, **p<0.01, ***p<0.001, ****p<0.0001. (B-G) Šidák multiple comparison post 2-way ANOVA; (H,J-M) Dunnett multiple comparison post 2-way ANOVA

There was no significant effect of cKD on body weight gain, fat mass or adiposity in APP/PS1 mice of either sex, though accretion of lean mass was greater in cKD-fed male mice (**Figs. 1C-F**, **S1A-D**). Despite minimal changes in weight, we observed KD-fed mice had dramatically increased caloric intake (**Fig. 1G**). As increased food intake in the absence of a difference in weight suggests altered energy balance, we placed mice in metabolic chambers; energy expenditure normalized to body weight was significantly increased in KD^on^ mice of both sexes (**Fig. 1H**). This difference was still seen when body weight was treated as a covariate, demonstrating that the greater energy expenditure of KD^on^-fed mice was not driven by body weight (**Fig. 1I**). RER was decreased in KD^on^ mice (**Fig. S1E**), indicating an increased reliance on fat oxidation for energy; this was expected as KD-fed mice do not consume any carbohydrates and rely primarily on fats for energy. Activity was not significantly altered in any of the groups (**Fig. S1F**).

To assess glucose tolerance and insulin sensitivity, we performed glucose and insulin tolerance tests. Glucose homeostasis was impaired in KD^on^ APP/PS1 mice, as indicated by increased fasting blood glucose (**Fig. S1G**), decreased glucose tolerance (**Figs. 1J-K**), and impaired insulin sensitivity (diet effect p=0.0018; **Figs. 1L-M**).

### Ketogenic diet improves cognition, amyloid load and mTOR activation in APP/PS1 mice

We performed a series of behavioral assays to assess the effect of KD on cognitive function and anxiety. There was no impact of diet on APP/PS1 performance in the open field test (**Fig. S2A**), but KD^on^ mice showed slightly greater anxiety in a dark/light box test as compared to KD^off^ mice (diet effect p=0.0310; **Fig. S2B**). In contrast, novel object recognition was improved substantially in female KD^on^ mice, with improved discrimination index and marginal improvements in absolute discrimination index (**Figs. 2A-B**), suggesting that female KD^on^ mice have both increased object discrimination and improved preference for the novel object. Spatial learning was not altered by diet during Barnes maze training (**Fig. 2C**), though this was most likely due to the overall good spatial memory of APP/PS1 mice. However, female KD-fed mice found the escape hole more quickly during the short-term memory test (**Fig. 2D**).

**Figure 2:**
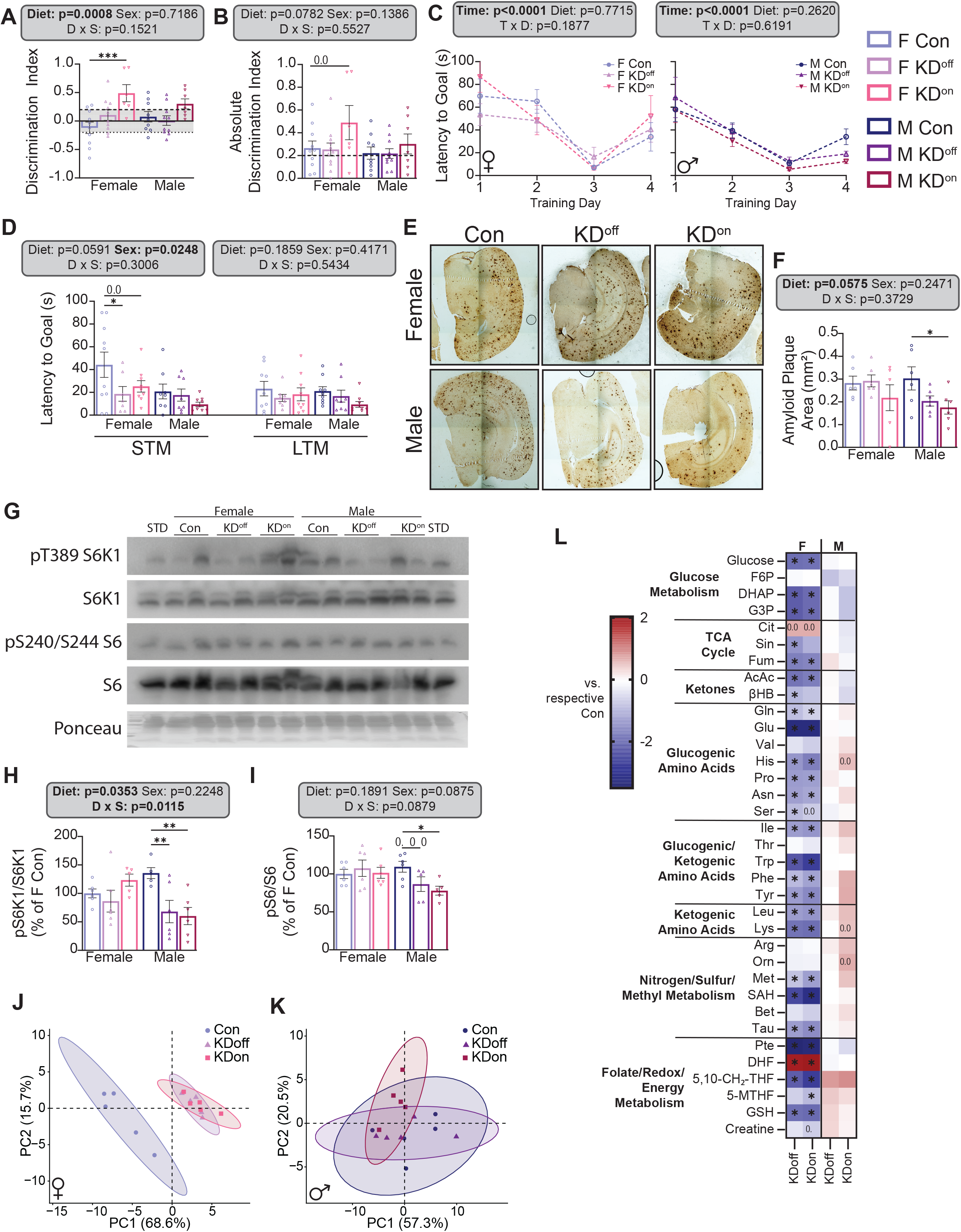
Ketogenic diet improves cognition, amyloid burden, mTOR signaling, and autophagy. A-B) Novel object recognition discrimination index (A) and absolute discrimination index (B). C-D) Barnes maze training period (C) and short- and long-term memory tests (D). E-F) DAB staining for 6E10 β-amyloid (E) and quantification of total plaque area (F). G-I) Immunoblotting for downstream targets of mTORC1 signaling (G). Quantification of pS6K/S6K (H) and pS6/S6 (I). J-K) Principle component analysis (PCA) of female (J) and male (K) brain polar metabolites. L) Brain polar metabolites. (A-D) Behavioral assays n=9-17/group; (E-N) histology, immunoblotting, and metabolomics n=4-6/group. Asterisks denote significant differences between diets: *p<0.05, **p<0.01, ***p<0.001, ****p<0.0001. (C) Šidák multiple comparison post 2-way ANOVA; (A-B, D-N) Dunnett multiple comparison post 2-way ANOVA

We also assessed the effect of KD on AD pathology. We found that cortical amyloid plaque area was reduced in mice on cKD, though this was more pronounced in males (**Figs. 2E-F**). Unlike in previous studies of KD, we observed no changes in microglial or astrocytic populations, or of aggregation around MX04-labeled amyloid plaques (**Figs. S2C-G**).

mTOR signaling and autophagy have been previously implicated in the pathogenesis of AD. We found that cKD (both in KD^off^ and KD^on^) reduced the phosphorylation of the mTORC1 substrate T389 S6K1 and the downstream mTORC1 readout S240/S244 S6 in APP/PS1 male but not female mice (**Figs. 2G-I**). Autophagy, in contrast, was not strongly altered by cKD (**Figs. S2H-L**).

We performed a metabolomics analysis of polar metabolites in whole brain to assess how cKD impacted aspects of brain metabolism. Principal component analysis (PCA) of mice fed Con or cKD diets revealed that APP/PS1 female mice exhibit a robust, long-lasting response to KD, with a high overlap between KD^off^ and KD^on^ mice (**Fig. 2J**). In contrast, APP/PS1 males had considerable overlap between all three diet groups (**Fig. 2K**). Analysis of metabolite levels revealed that a KD diet strongly alters metabolism in female mice, with the majority of changes seen during KD^on^ persisting into the KD^off^ period (**Fig. 2L**).

Surprisingly, levels of β-HB and acetoacetate were much lower in females (relative to Con) than in males and were similarly low in both KD^off^ and KD^on^ females as compared to Con (**Fig. 2L**). Finally, despite males having minimal changes in brain metabolites, metabolites involved in glucose metabolism (glucose, fructose-6-phosphate, dihydroxyacetone phosphate, and glyceraldehyde-3-phosphate) are reduced in both KD^off^ and KD^on^ compared to Con, suggesting that cKD may promote a shift away from glucose metabolism toward ketone metabolism.

### Ketogenic diet improves body weight and fat mass and acutely modifies whole-body metabolism and glucose homeostasis in 3xTg mice

We decided to explore the impact of cKD more deeply in the 3xTg mouse model of AD, which show age-related increases in amyloid and tau pathology, gliosis, and reliably show cognitive deficits in our lab (Oddo, Caccamo et al. 2003, Oddo, Caccamo et al. 2003, Babygirija, Sonsalla et al. 2024, Babygirija, Han et al. 2025, Babygirija, Green et al. 2026). 3xTg mice of both sexes were placed onto either Con or cKD diets at 6 months of age and followed until 13 months of age (**Fig. 3A**). Fasting ketone levels were significantly increased in KD^on^ males, but not females; however, after re-feeding ketone levels were significantly higher in both female and male KD^on^ mice (**Fig. 3B**).

**Figure 3:**
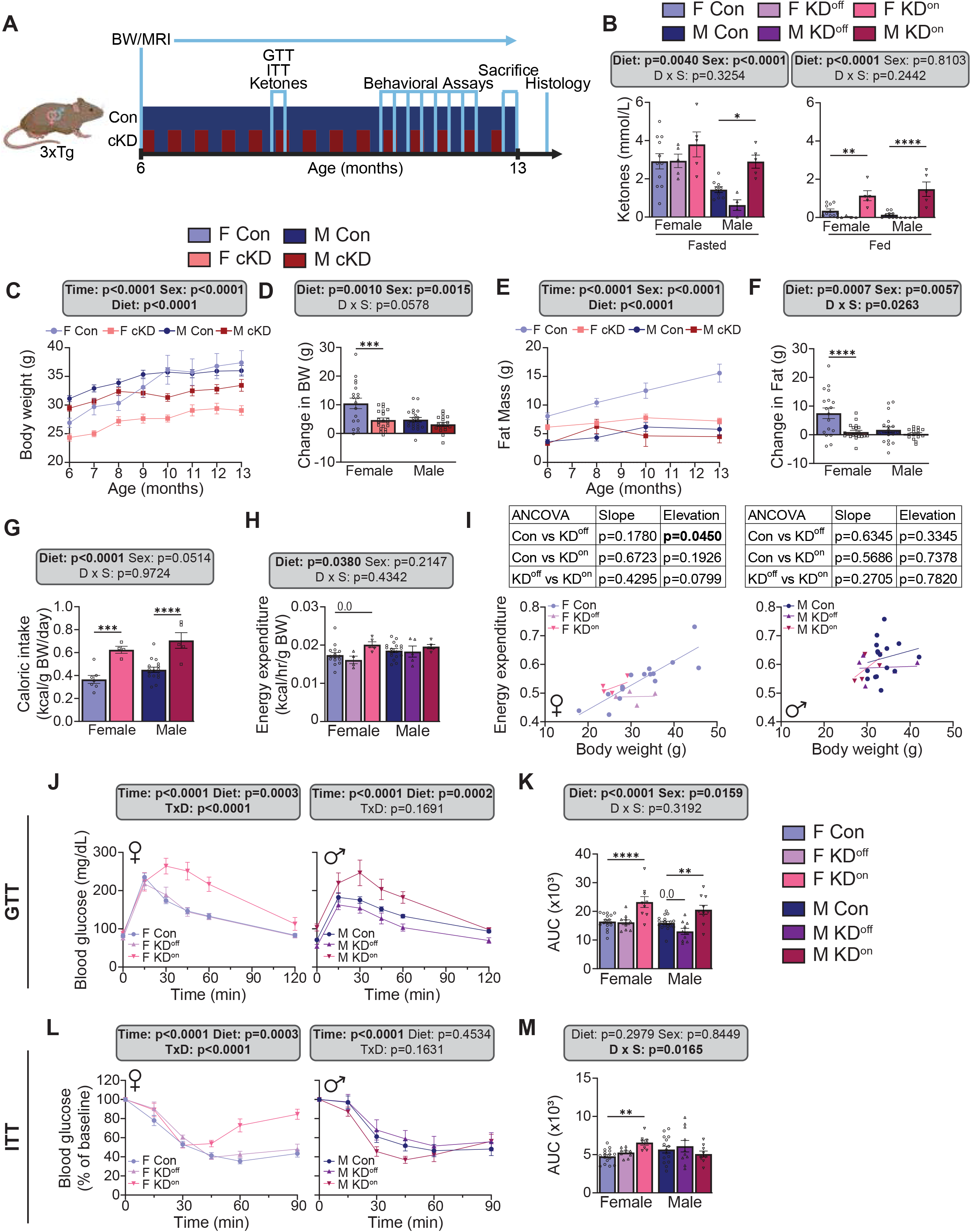
3xTg mice have improved body composition but worsened glucose homeostasis on ketogenic diet. A) Experiment Schematic B) Fasted and fed ketone levels C-D) Body weight across the experimental period (C) and change in body weight from 6 to 13 months (D). E-F) Fat mass across the experimental period (E) and change in fat mass from 6 to 13 months (F). G) Home cage food consumption normalized to body weight. H) Energy expenditure normalized to body weight. I) ANCOVA of energy expenditure. J-K) i.p. glucose tolerance test (J) and area under the curve (K). L-M) i.p. insulin tolerance test (L) and area under the curve (M). (A-F, H-M) Body weight/composition data n=18-20/group; (G) home cage data n=4-16/group; metabolic chamber data n=5-17/group; GTT/ITT data n=9-20/group;. Asterisks denote significant differences between diets: *p<0.05, **p<0.01, ***p<0.001, ****p<0.0001, (B-G) Šidák multiple comparison post 2-way ANOVA; (H,J-M) Dunnett multiple comparison post 2-way ANOVA

Female 3xTg mice fed cKD had reduced body weight and fat mass accretion compared to Con-fed mice, but diet did not affect final body weight or fat mass in males (**Figs. 3C-F**). Lean mass was not altered by diet in either sex (**Figs. S3A-B**). Adiposity was reduced by cKD in females, but not significantly affected in males (**Figs. S3C-D**). These improvements in female body composition occurred in the presence of significantly increased caloric intake in both sexes when on KD (**Fig. 3G**). This may be due to changes in energy expenditure, as mice on KD had increased energy expenditure when normalized to body weight (**Fig. 3H**). When energy expenditure was analyzed with weight as a covariate, this difference was revealed as primarily due to changes in body weight; however, there was a modest increase in KD^on^ females relative to KD^off^ female mice (p=0.0799); males did not have any significant changes to energy expenditure (**Fig. 3I**). RER was reduced in mice on KD as expected, while overall activity was not affected (**Figs. S3E-F**). Fasting blood glucose levels were increased in KD^on^ males, but unchanged by KD in females (**Fig. S3G**). KD^on^ mice of both sexes exhibited impaired glucose tolerance (**Figs. 3J-K**). KD^on^ females also exhibited impaired insulin sensitivity (**Figs. 3L-M**).

### Ketogenic diet acutely modifies spatial memory and AD pathology in 3xTg mice

We performed a series of behavioral assays to assess locomotor function, anxiety, and cognitive function. Neither female nor male mice exhibited significant alterations in movement during an open field test (**Fig. S4A**), anxiety during dark/light box assay (**Fig. S4B**), or changes in novel object recognition test (**Figs. S4C-D**). During Barnes maze, female KD^on^-fed mice showed improvements in spatial learning during training as evidenced by a reduction in latency to goal (**Fig. 4A**). While there were no changes to short-term memory in either sex, we observed an overall beneficial effect of diet (p=0.0308) on long-term memory in KD^on^ mice, which was statistically significant in male mice (**Fig. 4B**).

**Figure 4:**
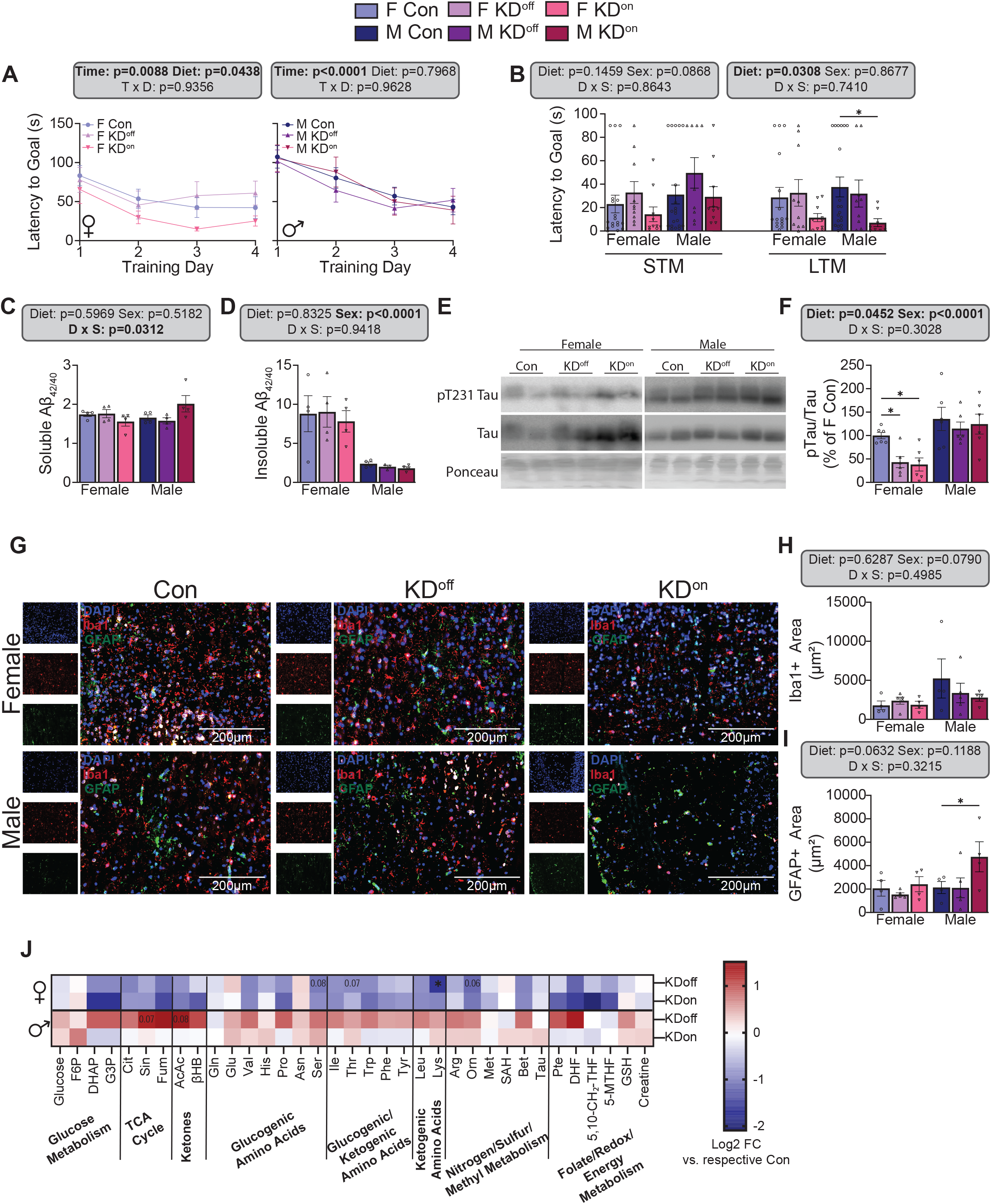
Ketogenic diet improves spatial memory and AD pathology in 3xTg mice. A-B) Barnes maze training period (A) and short- and long-term memory tests (B). C-D) Ratio of soluble (C) and insoluble (D) Aβ42/40. E-F) Immunoblotting of pT231 Tau and Tau (E) and quantification of pTau/Tau (F). G-I) Immunoflourescent staining of microglia (Iba1) and astrocytes (GFAP) (G) and quantification of total Iba1+ area (H) and GFAP+ area (I). J) Brain polar metabolites. (A-B) Barnes maze n=9-20/group. (C-N) Histology, immuoblotting and metabolomics n=4-6/group. Asterisks denote significant differences between diets: *p<0.05, **p<0.01, ***p<0.001, ****p<0.0001. (A) Šidák multiple comparison post 2-way ANOVA; (B-N) Dunnett multiple comparison post 2-way ANOVA

Although 3xTg mice have both amyloid and tau pathology, deposition of amyloid into plaques in this model is minor at 13 months of age (Sonsalla, Babygirija et al. 2025). We therefore performed an ELISA to measure the Aβ_42/40_ ratio to assess how cKD impacts amyloid pathology and found no changes in either sex (**Figs. 4C-D**). In contrast, the KD^on^ diet robustly reduced tau phosphorylation in female but not male 3xTg mice (**Figs. 4E-F**). We also observed an overall increase in astrocytic GFAP+ staining in male mice, while microglial staining was unchanged in both sexes (**Figs. 4G-I**). Finally, KD-fed 3xTg mice had decreased S6K1 T389 phosphorylation in both sexes (S6K1 diet effect p=0.0587), indicating mTORC1 inhibition (**Figs. S4E-G**), while autophagy was unchanged (**Figs. S4E, H-K**).

We found minor, sex-dependent changes in brain polar metabolites in response to KD. Unlike in APP/PS1 mice, there was significant overlap between all groups in PCA; this is likely due to large between subject variations within each diet group (**Figs. S4L-M**). Similar to APP/PS1, female and male mice show opposite responses in metabolite levels, with polar metabolites exhibiting an overall decrease in females and increase in males, though few metabolites were significantly altered (**Fig. 4J**).

### Sex and strain modify the impact of cKD on metabolic health, AD pathology, and cognition

We performed multivariate analysis on measures of metabolic health, cognition, and AD pathology to examine the role of strain in the response to cKD. During PCA, we found that strain strongly separates responses of both female and male mice in all diets (**Figs. 5A-B**). Interestingly, in female mice this separation was increased in KD^on^ mice but lessened in male KD^off^ and KD^on^ mice. This is likely driven by opposite responses to body composition in female mice. Indeed, upon examination of the contribution of each measure to PC1 and PC2, we found that body composition was a strong contributor to PC1-derived shifts (**Fig. S5A**). Visualizing all measures in a heatmap further revealed sex- and strain-dependent effects on the response to cKD, especially with regards to body composition, mTOR signaling, and autophagy (**Fig. 5C**).

**Figure 5:**
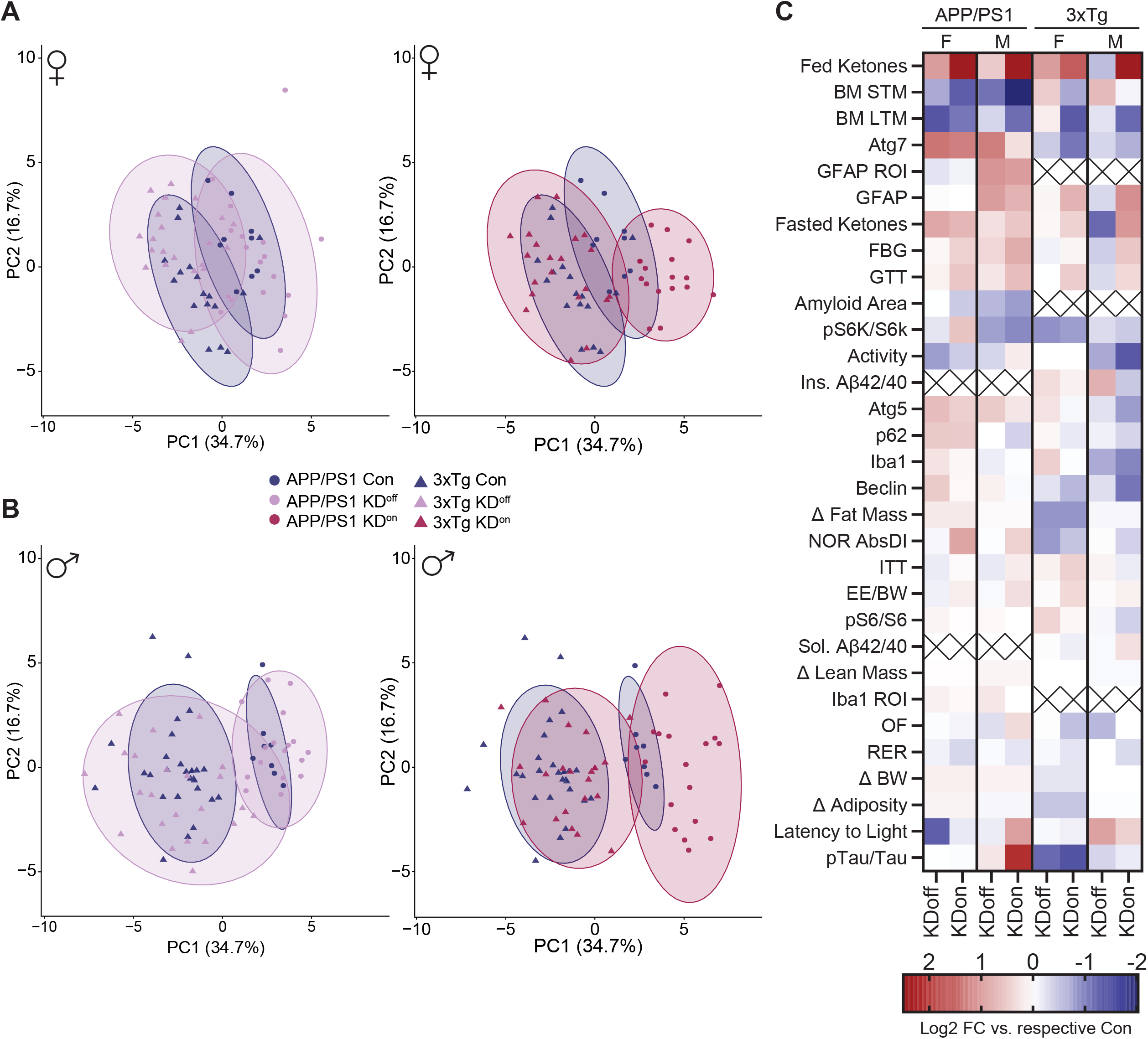
Sex and strain impact the response to cKD. A-B) Principle component analysis (PCA) of metabolic health, cognition, and AD pathology measures in female (A) and male (B) mice. C) Heat map of brain polar metabolite log2 fold changes compared to controls of each sex/strain. n=5/group.

### 1,3-butanediol supplementation worsens metabolic health in 3xTg mice

A KD is difficult to adhere to, which may limit its translatability. With this in mind, we examined the metabolic effects of two dietary supplements, 1,3-butanediol (1,3-BD) and medium chain triglycerides (MCT), which have been previously shown to increase ketone levels in mice, and as such may be promising KD-mimicking interventions.

We supplemented 3xTg mice with either 1,3-butanediol (1,3-BD) or medium chain triglycerides (caprylic acid; MCT) in the diet from 6 to 13 months of age (**Fig. 6A**). During fasting, female mice supplemented with MCT had increased ketone levels; during the fed period 1,3-BD supplementation increased ketones in both sexes (**Fig. 6B**). 1,3-BD supplementation increased weight and fat mass gain in both sexes (**Figs. 6C-F**), increasing adiposity without altering lean mass (**Figs. S6A-D**). These changes occurred in the absence of altered food consumption or energy expenditure normalized to body weight (**Figs. 6G-H**). However, when body weight was analyzed as a covariate, we identified a paradoxical increase in energy expenditure in 1,3-BD-fed female mice as compared to Con, but not in males (**Fig. 6I**). RER and activity were also not altered by diet (**Figs. S6E-F**). Glucose homeostasis was impaired in 1,3-BD-fed mice, indicated by increased fasting blood glucose levels in females (**Fig. S6G**) and decreased glucose tolerance (**Figs. 6J-K**) in both sexes; insulin sensitivity was impaired by 1,3-BD in both sexes but reached significance only in male mice (**Figs. 6L-M**).

**Figure 6:**
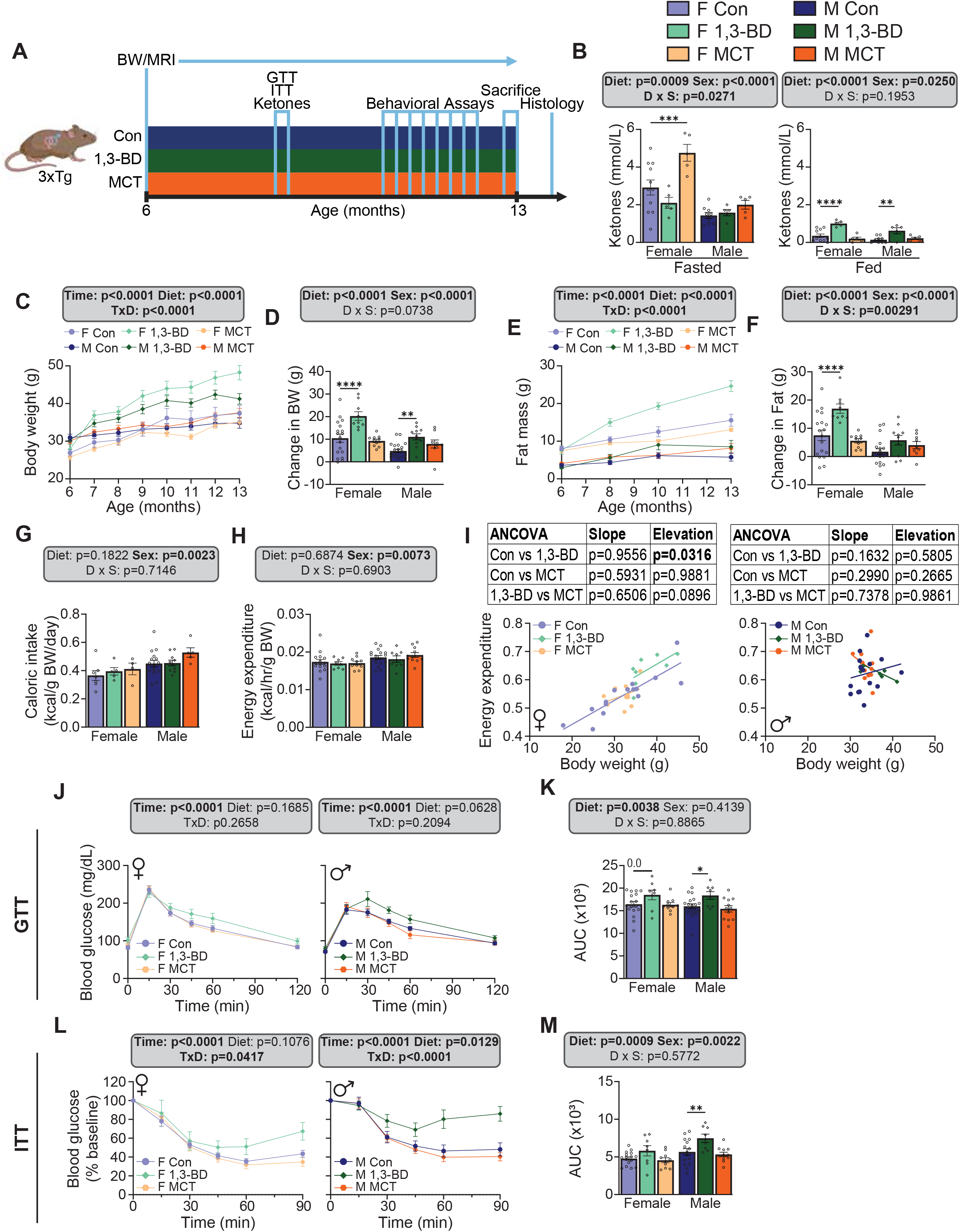
1,3-BD worsens metabolic health in 3xTg mice. A) Experiment Schematic B) Fasted and fed ketone levels C-D) Body weight across the experimental period (C) and change in body weight from 6 to 13 months (D). E-F) Fat mass across the experimental period (E) and change in fat mass from 6 to 13 months (F). G) Home cage food consumption normalized to body weight. H) Energy expenditure normalized to body weight. I) ANCOVA of energy expenditure. J-K) i.p. glucose tolerance test (J) and area under the curve (K). L-M) i.p. insulin tolerance test (L) and area under the curve (M). (A-F, H-M) Body weight/composition, metabolic chambers, and GTT/ITT data n=7-20/group; (G) home cage data n=4-16/group. Asterisks denote significant differences between diets: *p<0.05, **p<0.01, ***p<0.001, ****p<0.0001. (B-G) Šidák multiple comparison post 2-way ANOVA; (H,J-M) Dunnett multiple comparison post 2-way ANOVA.

### 1,3-butanediol supplementation worsens locomotion during open field test but improves object recognition in 3xTg mice

We found that 1,3-BD-fed females had decreased movement in open field test, suggesting a possible increase in anxiety, but no effect on behavior in the dark/light box test was observed (**Figs. S7A-B**). In contrast, novel object recognition performance was improved in 1,3-BD-fed mice (**Figs. 7A-B**). Discrimination index was positively elevated in 1,3-BD mice in both sexes (**Fig. 7A**), while absolute discrimination index was unchanged (**Fig. 7B**), suggesting that changes in discrimination index are driven by improved preference for the novel object which may represent decreased neophobia. During the 4-day Barnes maze training period, female mice on 1,3-BD and MCT had minor impairments to spatial learning (diet effect: p=0.0647) while male mice had a significant diet effect (p=0.0299) which we interpret as a slight improvement in MCT-fed mice and slight worsening in 1,3-BD-fed mice (**Fig. 7C**). No differences were seen during Barnes maze short- or long-term memory tests (**Fig. 7D**).

**Figure 7:**
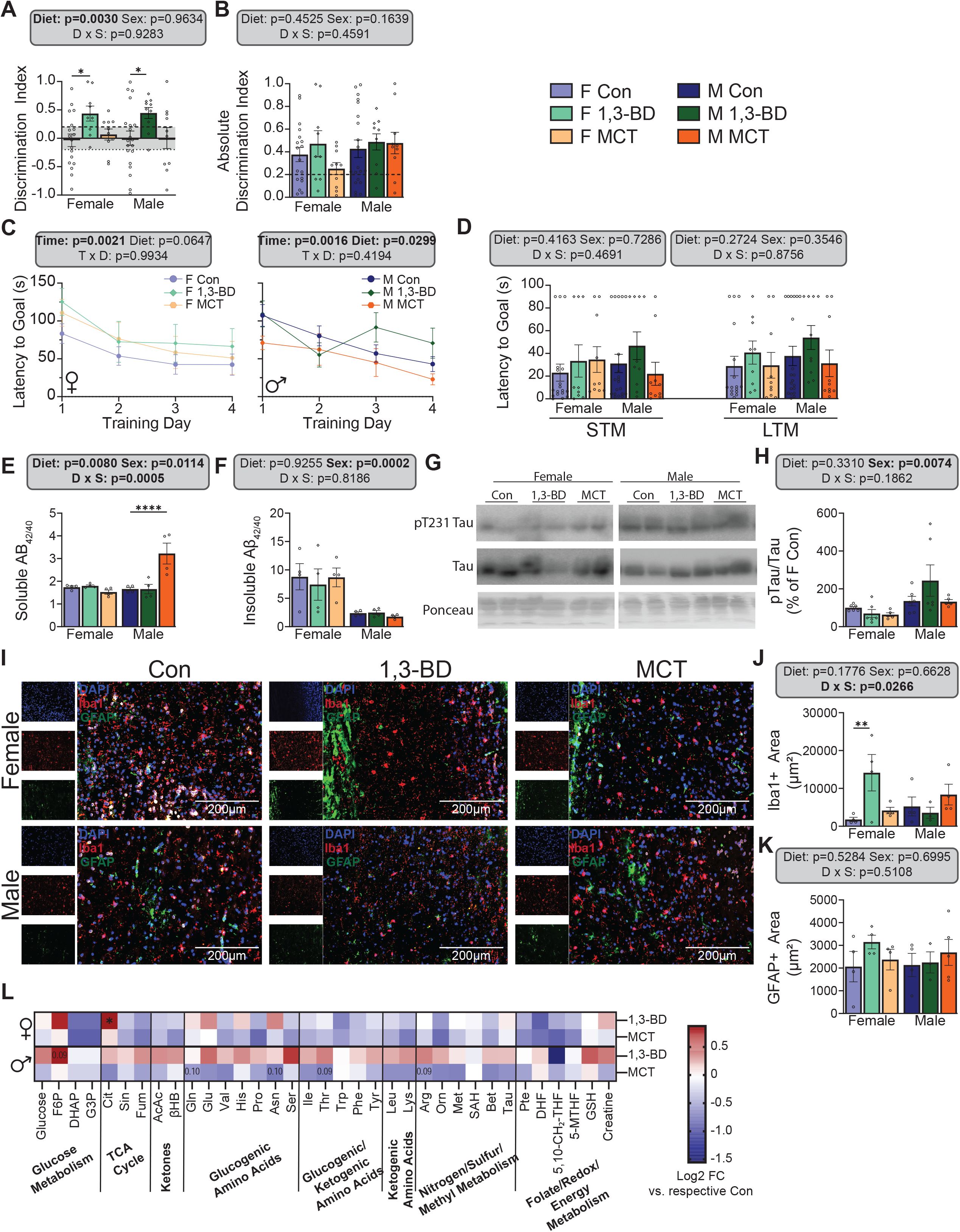
1,3-BD improves object recognition. A-B) Novel object recognition discrimination index (A) and absolute discrimination index (B) C-D) Barnes maze training period (C) and short- and long-term memory tests (D). E-F) Ratio of soluble (E) and insoluble (F) Aβ42/40. G-H) Immunoblotting of pT231 Tau and Tau (G) and quantification of pTau/Tau (H). I-K) Immunoflourescent staining of microglia (Iba1) and astrocytes (GFAP) (I) and quantification of total Iba1+ area (J) and GFAP+ area (K). L) Heat map of brain polar metabolites. (A-D) Behavioral assays n=7-20/group. (E-K) Histology and metabolomics n=4-6/group. Asterisks denote significant differences between diets: *p<0.05, **p<0.01, ***p<0.001, ****p<0.0001. (C) Šidák multiple comparison post 2-way ANOVA; (A-B, D-K) Dunnett multiple comparison post 2-way ANOVA

Next, we assessed the role of ketone supplementation on AD pathology and molecular signaling. We found that MCT increased soluble Aβ_42/40_ in male 3xTg mice, with no effect of MCT in females or of 1,3-BD in either sex; neither compound had an effect on tau phosphorylation (**Figs. 7E-H**). Cortical Iba1+ microglial staining was increased by 1,3-BD in female, but not male mice, while MCT had no effect; astrocytes were not altered by either compound (**Figs. 7I-K**). Finally, we observed decreased S6 phosphorylation in both 1,3-BD and MCT-treated males, suggesting a sex-specific decrease in mTORC1 activity (**Figs. S7C-I**). In 3xTg mice fed diets containing either 1,3-BD or MCT, PCA revealed no major shifts in metabolites in females or males (**Figs. S7J-K**). Neither sex exhibited strong metabolite changes in response to either supplement, though males exhibited decreases in select amino acids in response to MCT (**Fig. 7L**).

## Discussion

There is high interest in identifying dietary interventions that can combat AD pathology and cognitive deficits. Here, we examined the effect of a cyclic ketogenic diet (KD) on metabolic health, cognition, and AD pathology in two AD mouse models. cKD increased fed ketone levels across sexes and strains, but impaired glycemic control in all groups. In contrast, the effects of cKD on body composition are both sex- and strain-dependent, as body composition is worsened in APP/PS1 females and improved in 3xTg females fed cKD. Despite these negative effects on metabolic health, cKD reliably improved several measures of cognition and AD pathology; these benefits were also sex- and strain-dependent. In APP/PS1 mice, cognitive benefits were stronger in females, while the effects on amyloid levels, mTOR, and autophagy were greater in males. In 3xTg mice, improvements in spatial memory were associated with improved amyloid pathology in males and tau pathology in females.

We additionally examined the impact of two ketone supplements, 1,3-BD and MCT, in the 3xTg mouse model of AD. 1,3-BD and MCT both result in elevated ketone levels under either fed or fasted conditions, but they have very different metabolic and AD responses compared to cKD-fed mice. MCT treatment did not impact metabolic health, cognition, or AD pathology. In contrast, 1,3-BD improved cognition, but did not alter AD pathology and had strongly negative effects on metabolic health, particularly in females. One possible reason for the reduced benefits of these supplements as compared to a cKD is that the treatments did not elevate ketones as effectively as a cKD. Alternatively, elevated ketones alone may be insufficient to achieve the full benefits of a cKD; perhaps ketogenesis itself has beneficial effects through reduced carbohydrate availability or increases in fatty acid metabolism.

cKD worsened glucose homeostasis across sex and strain, but only when mice were actively on KD (KD^on^); while the cKD mice were consuming the Con diet, mice returned to baseline. The impaired glycemic control when consuming the KD diet is likely the result of the metabolic shifts induced by the adaptation to extreme carbohydrate restriction, and has been previously reported (Pehleman, Peters et al. 2005, Webster, van Boom et al. 2020). Importantly, these effects are rapidly reversable, with negative effects on glucose tolerance and insulin sensitivity ameliorated within 5 days of returning to a standard diet (Kinzig, Honors et al. 2010). Together with our observation that glucose intolerance was present only during KD^on^ weeks and was abolished following reintroduction of the Con diet, these findings support the possibility that this phenotype represents a dynamic adaptation to carbohydrate restriction rather than a persistent deterioration in metabolic health. We find, as have others, that these effects are temporary and reverse following the return to a normal diet.

Despite the rapidly reversible metabolic adaptations, we found that alterations in brain metabolites were more stable, with similar levels in both KD^on^ and KD^off^ mice compared to Con. This suggests that mice may maintain a tighter regulation of brain metabolite levels. In addition, brain levels of acetoacetate and β-HB were decreased or unchanged despite increased plasma levels in KD^on^, 1,3-BD, and MCT-fed mice, and previous work in both rodents (Hawkins, Mans et al. 1986, Bentourkia, Tremblay et al. 2009) and humans (Owen, Morgan et al. 1967, Hasselbalch, Knudsen et al. 1995, Blomqvist, Alvarsson et al. 2002) have shown that brain ketones are directly proportional to systemic levels, we would expect brain levels of acetoacetate (AcAc) and β-HB to be likewise increased. The lack of increase in brain ketone levels may therefore be due to increased ketone utilization and further emphasizes differences in systemic and brain metabolism.

We found that elevation of systemic ketone levels, through diet or supplementation, was generally associated with improved cognition. These findings are consistent with previous preclinical and clinical studies reporting cognitive benefits following ketogenic diets or ketone supplementation, although the magnitude and domains of benefit vary substantially between studies (Kashiwaya, Bergman et al. 2013, Grammatikopoulou, Goulis et al. 2020, Xu, Jiang et al. 2022, Rong, Peng et al. 2024, Abdel-Aziz, Ahmed et al. 2025, Chen, Li et al. 2025, Di Lucente, Rutkowsky et al. 2026, Fulghum, Hayir et al. 2026). Here, we show that improvements to cognition in cKD-fed mice are sex and strain dependent, with cKD improving novel object recognition and spatial memory in female, but not male APP/PS1 mice, and improving spatial memory, but not novel object recognition, in both sexes of 3xTg mice. In addition to sex and strain effects, there are also important differences in the timing of the benefits of the cKD. Cognition was primarily improved only during active consumption of the KD (KD^on^), and were not observed during KD^off^; thus, these improvements are likely to acute cKD-induced changes, e.g. in ketone signaling or fuel utilization, rather than engaging longer-term mechanisms such as tau or amyloid clearance.

Potential limitations of the current work include the use of mouse models in studying KD. It has been previously established that mice and humans respond differently to KD with regards to metabolic health. KD in mice is strongly associated with hyperlipidemia and worsened glucose homeostasis (Gallop, Vieira et al. 2025), as we see here, but in humans KD has been shown to improve body weight and glucose homeostasis in obese and diabetic patients (Dowis and Banga 2021). Utilizing a cKD paradigm has been previously shown to be effective at reducing the metabolic dysfunction associated with KD in mice (Newman, Covarrubias et al. 2017), but we find that this is a strain-dependent response. Additionally, AD mouse models are more representative of early onset AD as opposed to sporadic AD, and as such may not effectively represent all aspects of human AD pathology. While we did study the impact of cKD in two different mouse models, differences between the models complicate our ability to draw strong conclusions. Genetic variations between APP/PS1 and 3xTg include different background strains (C57BL/6JxC3H vs B6129SF2/J, respectively) and different AD-related mutations (APP695_Swe_/PS1-dE9 and APP_Swe_/Tau_P301L_/Psen1, respectively), resulting in different aspects and trajectories of AD pathology. We chose to start our experiments at slightly different ages due to this to capture the point in each model when pathology has already begun but before substantial cognitive deficits could occur. Future work should try to further dissect if the strain-dependent differences are due to background genetics or pathological features. Finally, we examined glial populations, mTORC1 activity, and autophagy as possible mechanisms that may be involved in the improvements to AD pathology and cognition that we see. While S6K phosphorylation was reduced in male APP/PS1 mice and there was an overall diet effect (p=0.0587) in 3xTg mice, we did not see robust changes to mTORC1 signaling, nor did we see much alterations in glial populations or autophagy markers. Future work should further investigate the potential role of mTORC1 but also examine changes to brain metabolism including mitochondrial function. The transient nature of the benefits of KD that we see here suggest that improvements to cognition may be mediated by the ketones themselves.

This work has important implications for the treatment of AD. Not only do we show that ketones are capable of improving AD pathology and cognition independent of improvements in metabolic health, we also reveal that sex and genetic background impact the response. Our 3xTg mice exhibited sexually dimorphic effects on AD pathology, converging on improvements in cognition in both sexes. APP/PS1 mice had more consistent responses in amyloid pathology, but only female mice had improved novel object recognition. We showed that while KD is a promising intervention, it may be associated with negative metabolic consequences. Supplements that increase ketone levels may be an effective option for AD, but more research needs to be done to find ketone supplements that are more effective at recapitulating the benefits of KD without negative metabolic consequences. These findings are important due to the high genetic variability in human populations and the differences in AD pathology/cognition between men and women; limitations of current pharmaceutical AD treatments may have limited benefits due to these between-subject differences.

## Data Availability Statement

The data that support the plots within this article and other findings of this study, including full scans of western blot images, are provided as Source Data files

## Acknowledgements

We would like to thank all members of the Lamming lab for their assistance and input. The Lamming laboratory is supported in part by the NIH/NIA (AG056771, AG081482, AG084156, AG085898 and AG094153 to D.W.L.), a grant from the Alzheimer’s Association (23AARG-1029665), and startup funds from the University of Wisconsin-Madison School of Medicine and Public Health and Department of Medicine to D.W.L. C-Y.Y. was supported in part by a NIA K99/R00 award (K99AG084921). R.B. was supported by F31AG081115. RB is a UW Distinguished Research Fellow; support for this research was provided by the University of Wisconsin-Madison Office of the Vice Chancellor for Research with funding from the Wisconsin Alumni Research Foundation. The Puglielli laboratory is supported by the National Institutes of Health (R01NS094154, R01GM148487, and R01AG078794). The authors used the UW Carbone Cancer Center Experimental Animal Pathology Laboratory (P30CA014520) for tissue sectioning and histology. S.N.C., D.A.H, L.P., and D.W.L. are members of the Wisconsin Nathan Shock Center of Excellence in the Basic Biology of Aging, P30 AG092586. The Lamming lab was supported in part by the U.S. Department of Veterans Affairs (IS1-BX005524), and this work was supported using facilities and resources from the William S. Middleton Memorial Veterans Hospital. A total of approximately $237,500, or 95% percent of this project was financed with federal funds. A total of approximately $12,500, or 5% percent of this project was financed with non-governmental funds. The content is solely the responsibility of the authors and does not necessarily represent the official views of the NIH. This work does not represent the views of the Department of Veterans Affairs or the United States Government.

