## Supplemental Figures and Table Legends for "Ketones uncouple Alzheimer’s disease pathology and cognition from metabolic health in two mouse models"

Figure S1

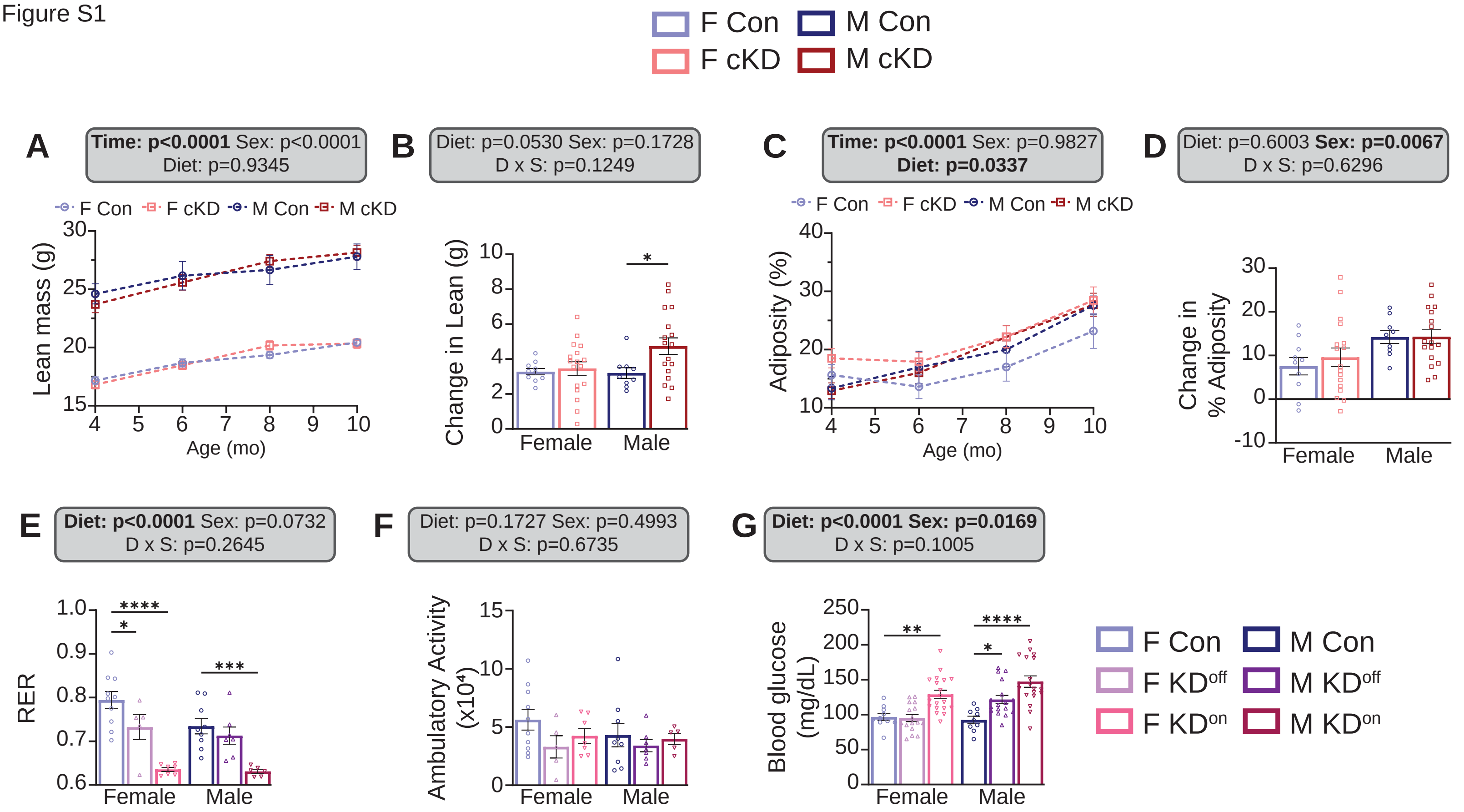
F Con
M Con
F cKD
M cKD
A
Time: p<0.0001 Sex: p<0.0001
Diet: p=0.9345
Lean mass (g)
Age (mo)
B
Diet: p=0.0530 Sex: p=0.1728
D x S: p=0.1249
Change in Lean (g)
Female
Male
C
Time: p<0.0001 Sex: p=0.9827
Diet: p=0.0337
Adiposity (%)
Age (mo)
D
Diet: p=0.6003 Sex: p=0.0067
D x S: p=0.6296
Change in % Adiposity
E
Diet: p<0.0001 Sex: p=0.0732
D x S: p=0.2645
RER
F
Diet: p=0.1727 Sex: p=0.4993
D x S: p=0.6735
Ambulatory Activity (x10⁴)
G
Diet: p<0.0001 Sex: p=0.0169
D x S: p=0.1005
Blood glucose (mg/dL)
F KDoff
M KDoff
F KDon
M KDon

**Figure S1: Ketogenic diet reduces RER and increases fasting blood glucose in APP/PS1 mice.**

A-B) Lean mass across the experimental period (A) and change in lean mass from 4 to 10 months (B).

C-D) Adiposity across the experimental period (C) and change in adiposity from 4 to 10 months (D).

E-F) RER (E) and ambulatory activity (F) during metabolic chambers.

G) Fasting blood glucose after 16hr overnight fast.

(A-G) Body weight/composition, metabolic chamber, and FBG data n=9-17/group. Asterisks denote significant differences between diets: *$p<0.05$, **$p<0.01$, ***$p<0.001$, ****$p<0.0001$. (A-D) Šidák multiple comparison post 2-way ANOVA; (E-G) Dunnett multiple comparison post 2-way ANOVA.

Figure S2

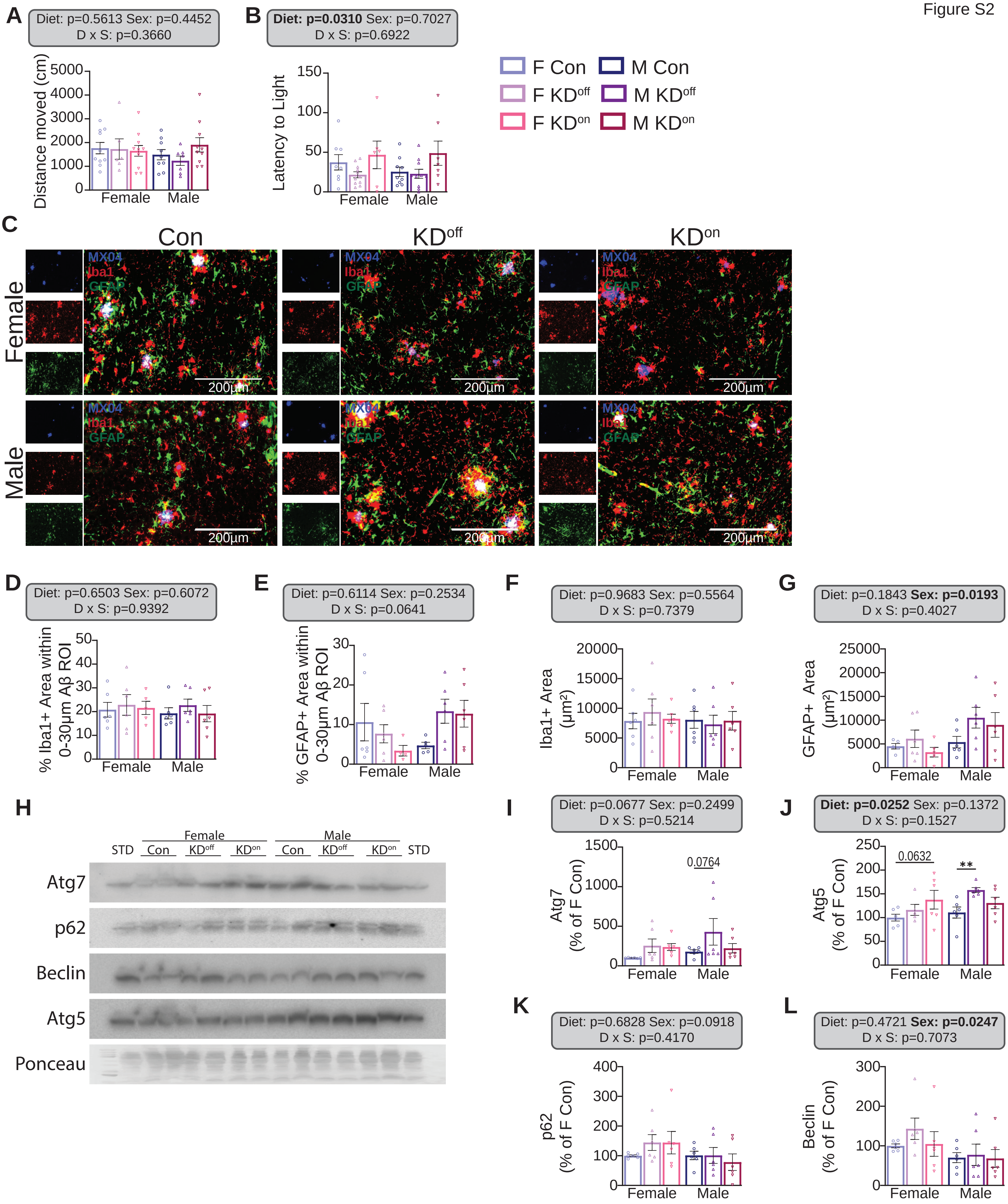

A
Diet: p=0.5613 Sex: p=0.4452
D x S: p=0.3660
Distance moved (cm)
Female
Male
B
Diet: p=0.0310 Sex: p=0.7027
D x S: p=0.6922
Latency to Light
F Con
M Con
F KDoff
M KDoff
F KDon
M KDon
C
Con
KDoff
KDon
Female
Male
MX04
Iba1
GFAP
200µm
D
Diet: p=0.6503 Sex: p=0.6072
D x S: p=0.9392
% Iba1+ Area within 0-30µm Aβ ROI
E
Diet: p=0.6114 Sex: p=0.2534
D x S: p=0.0641
% GFAP+ Area within 0-30µm Aβ ROI
F
Diet: p=0.9683 Sex: p=0.5564
D x S: p=0.7379
Iba1+ Area (µm²)
G
Diet: p=0.1843 Sex: p=0.0193
D x S: p=0.4027
GFAP+ Area (µm²)
H
STD
Con
KDoff
KDon
Atg7
p62
Beclin
Atg5
Ponceau
I
Diet: p=0.0677 Sex: p=0.2499
D x S: p=0.5214
Atg7 (% of F Con)
0.0764
J
Diet: p=0.0252 Sex: p=0.1372
D x S: p=0.1527
Atg5 (% of F Con)
0.0632
**
K
Diet: p=0.6828 Sex: p=0.0918
D x S: p=0.4170
p62 (% of F Con)
L
Diet: p=0.4721 Sex: p=0.0247
D x S: p=0.7073
Beclin (% of F Con)

**Figure S2: Ketogenic diet worsens anxiety in dark/light box assay.**
A) Distance moved in open field test.
B) Latency to light portion when placed in dark portion of dark/light box.
C-G) Immunoflourescent staining for amyloid (MethoxyX04), microglia (Iba1), and astrocytes (GFAP) (C) and quantified % positive areas of Iba1 (D) and GFAP (E) within a 0-30 μm region of interest around MX04-positive areas. Quantified total Iba1+ area (F) and GFAP+ area (G).
H-L) Immunoblotting for autophagy markers (H). Quantification of Atg7 (I; normalized to ponceau), Atg5 (J; normalized to ponceau), p62 (K; normalized to ponceau) and Beclin (L; normalized to ponceau).
(A-D) Behavioral assays n=9-17/group; (C-L) histology, immunoblotting, metabolomics n=4-6/group. Asterisks denote significant differences between diets: *$p<0.05$, **$p<0.01$, ***$p<0.001$, ****$p<0.0001$. (A-G) Dunnett multiple comparison post 2-way ANOVA

Figure S3

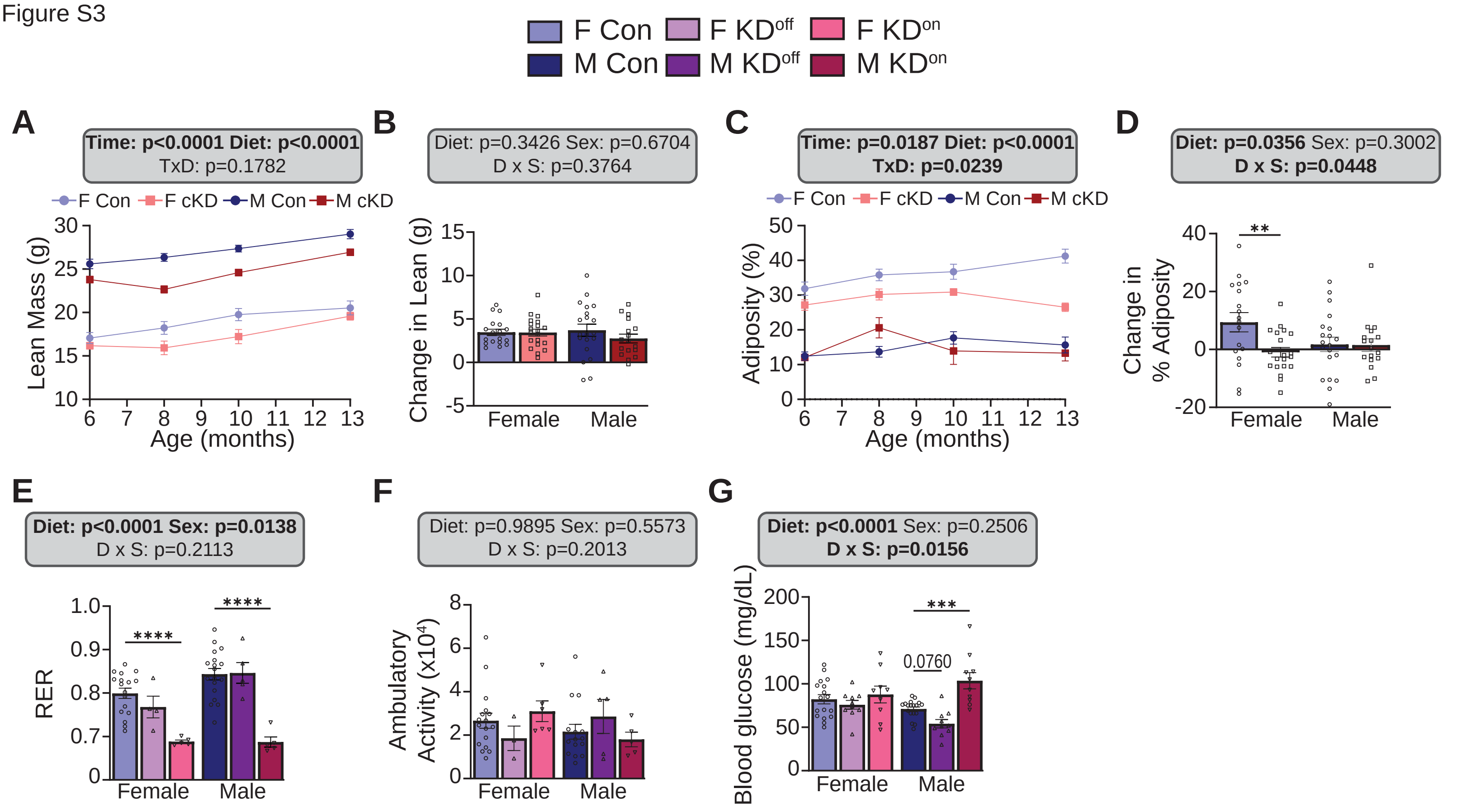
F Con
F KDoff
F KDon
M Con
M KDoff
M KDon
A
Time: p<0.0001 Diet: p<0.0001
TxD: p=0.1782
F Con
F cKD
M Con
M cKD
Lean Mass (g)
Age (months)
B
Diet: p=0.3426 Sex: p=0.6704
D x S: p=0.3764
Change in Lean (g)
Female
Male
C
Time: p=0.0187 Diet: p<0.0001
TxD: p=0.0239
Adiposity (%)
Age (months)
D
Diet: p=0.0356 Sex: p=0.3002
D x S: p=0.0448
Change in
% Adiposity
**
E
Diet: p<0.0001 Sex: p=0.0138
D x S: p=0.2113
RER
****
F
Diet: p=0.9895 Sex: p=0.5573
D x S: p=0.2013
Ambulatory
Activity (x10^4)
G
Diet: p<0.0001 Sex: p=0.2506
D x S: p=0.0156
Blood glucose (mg/dL)
0.0760
***

**Figure S3: Ketogenic diet improves adiposity and lowers RER.**

A-B) Lean mass across the experimental period (A) and change in lean mass from 6 to 13 months (B).

C-D) Fat mass across the experimental period (C) and change in fat mass from 6 to 13 months (D).

E-F) RER (E) and ambulatory activity (F) during metabolic chambers.

G) Fasting blood glucose after 16hr overnight fast.

(A-D) Body weight/composition data n=18-20/group; (E-F) metabolic chamber data n=5-17/group; (G) FBG data n=9-20/group;. Asterisks denote significant differences between diets: *$p<0.05$, **$p<0.01$, ***$p<0.001$, ****$p<0.0001$. (A-D) Šidák multiple comparison post 2-way ANOVA; (E-G) Dunnett multiple comparison post 2-way ANOVA

Figure S4

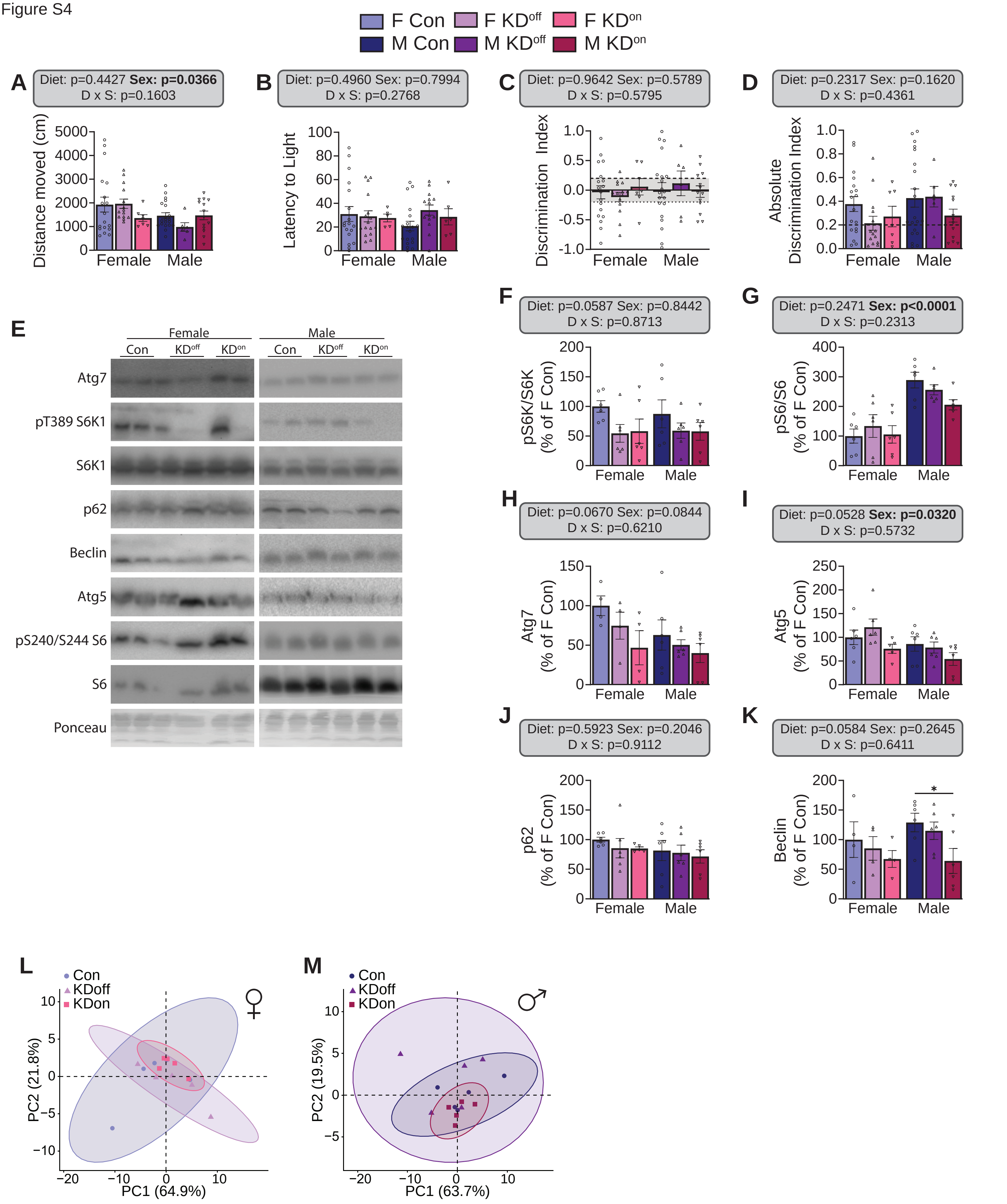

F Con
F KDoff
F KDon
M Con
M KDoff
M KDon
A
Diet: p=0.4427 **Sex: p=0.0366**
D x S: p=0.1603
Distance moved (cm)
Female
Male
B
Diet: p=0.4960 Sex: p=0.7994
D x S: p=0.2768
Latency to Light
C
Diet: p=0.9642 Sex: p=0.5789
D x S: p=0.5795
Discrimination Index
D
Diet: p=0.2317 Sex: p=0.1620
D x S: p=0.4361
Absolute Discrimination Index
E
Female
Male
Con
KDoff
KDon
Atg7
pT389 S6K1
S6K1
p62
Beclin
Atg5
pS240/S244 S6
S6
Ponceau
F
Diet: p=0.0587 Sex: p=0.8442
D x S: p=0.8713
pS6K/S6K (% of F Con)
G
Diet: p=0.2471 **Sex: p<0.0001**
D x S: p=0.2313
pS6/S6 (% of F Con)
H
Diet: p=0.0670 Sex: p=0.0844
D x S: p=0.6210
Atg7 (% of F Con)
I
Diet: p=0.0528 **Sex: p=0.0320**
D x S: p=0.5732
Atg5 (% of F Con)
J
Diet: p=0.5923 Sex: p=0.2046
D x S: p=0.9112
p62 (% of F Con)
K
Diet: p=0.0584 Sex: p=0.2645
D x S: p=0.6411
Beclin (% of F Con)
*
L
Con
KDoff
KDon
PC1 (64.9%)
PC2 (21.8%)
M
Con
KDoff
KDon
PC1 (63.7%)
PC2 (19.5%)

**Figure S4: Behavioral assays and autophagy markers in cKD-fed 3xTg mice.**

A) Distance moved in open field test.

B) Latency to light portion when placed in dark portion of dark/light box.

C-D) Novel object recognition discrimination index (C) and absolute discrimination index (D)

E-K) Immunoblotting for downstream targets of mTORC1 and autophagy markers (E). Quantification of pS6K/S6K (F), pS6/S6 (G), Atg7 (H; normalized to ponceau), Atg5 (I; normalized to ponceau), p62 (J; normalized to ponceau) and Beclin (K; normalized to ponceau).

L-M) PCA of brain polar metabolites in female (L) and male (M) mice.

(A-D) Behavioral assays n=9-20/group. (E-M) Immunoblotting and metabolomics n=4-6/group. Asterisks denote significant differences between diets: *$p<0.05$, **$p<0.01$, ***$p<0.001$, ****$p<0.0001$. (A-I) Dunnett multiple comparison post 2-way ANOVA

Figure S5

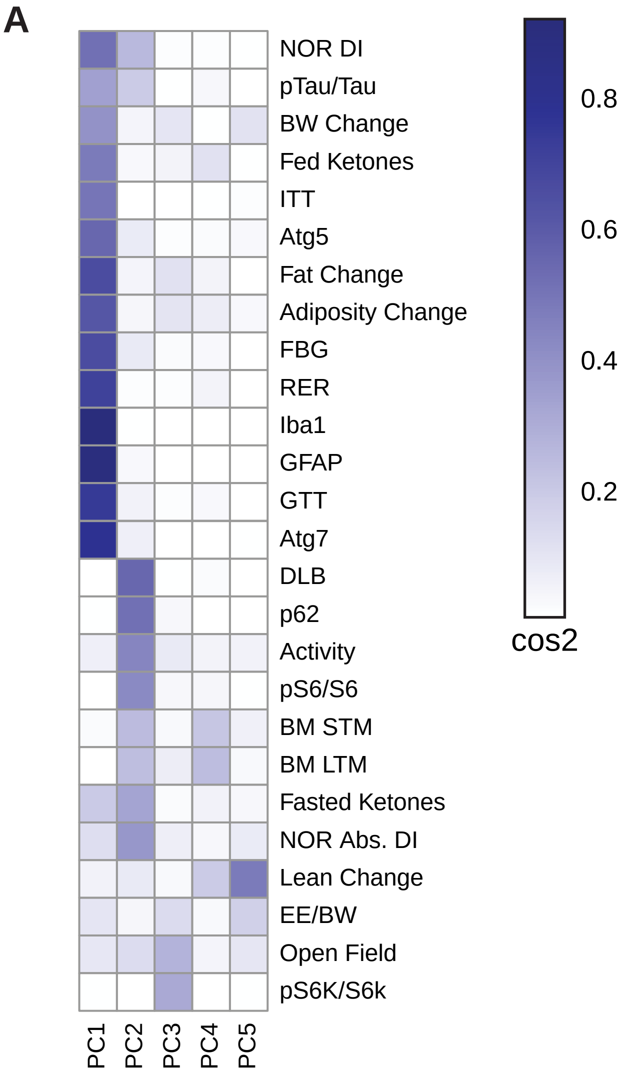
A
NOR DI
pTau/Tau
BW Change
Fed Ketones
ITT
Atg5
Fat Change
Adiposity Change
FBG
RER
Iba1
GFAP
GTT
Atg7
DLB
p62
Activity
pS6/S6
BM STM
BM LTM
Fasted Ketones
NOR Abs. DI
Lean Change
EE/BW
Open Field
pS6K/S6k
PC1
PC2
PC3
PC4
PC5
0.8
0.6
0.4
0.2
cos2

**S5: Contribution of measures to PCA dimensions.**

A) Contribution of measures to PCAs in 5A-B.

Figure S6

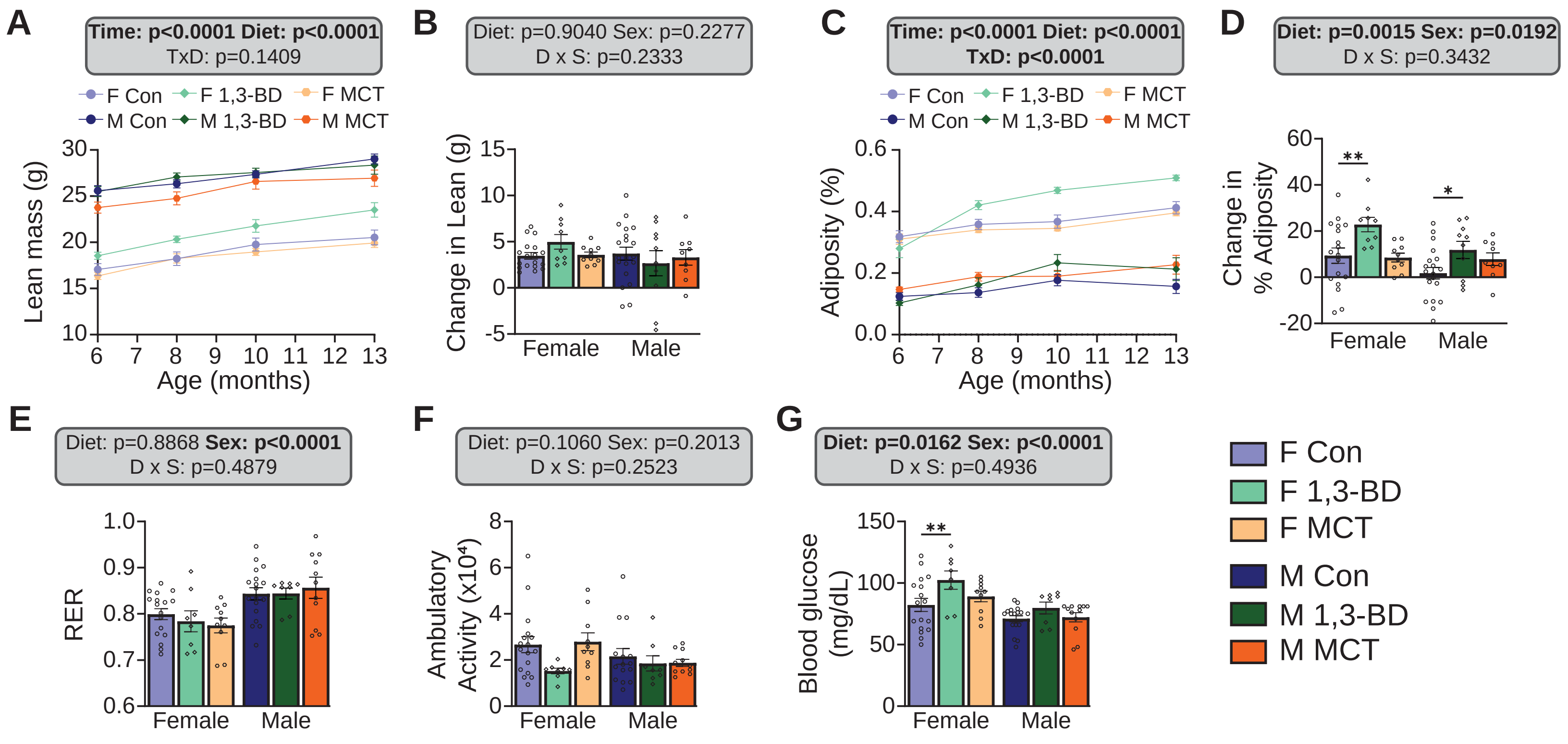

A
Time: p<0.0001 Diet: p<0.0001
TxD: p=0.1409
F Con
F 1,3-BD
F MCT
M Con
M 1,3-BD
M MCT
Lean mass (g)
Age (months)
B
Diet: p=0.9040 Sex: p=0.2277
D x S: p=0.2333
Change in Lean (g)
Female
Male
C
Time: p<0.0001 Diet: p<0.0001
TxD: p<0.0001
Adiposity (%)
Age (months)
D
Diet: p=0.0015 Sex: p=0.0192
D x S: p=0.3432
Change in % Adiposity
**
*
Female
Male
E
Diet: p=0.8868 Sex: p<0.0001
D x S: p=0.4879
RER
Female
Male
F
Diet: p=0.1060 Sex: p=0.2013
D x S: p=0.2523
Ambulatory Activity (x10⁴)
Female
Male
G
Diet: p=0.0162 Sex: p<0.0001
D x S: p=0.4936
Blood glucose (mg/dL)
**
Female
Male
F Con
F 1,3-BD
F MCT
M Con
M 1,3-BD
M MCT

**Figure S6: 1,3-BD worsens metabolic health in 3xTg mice.**

A-B) Lean mass across the experimental period (A) and change in lean mass from 6 to 13 months (B).

C-D) Fat mass across the experimental period (C) and change in fat mass from 6 to 13 months (D).

E-F) RER (E) and ambulatory activity (F) during metabolic chambers.

G) Fasting blood glucose after 16hr overnight fast.

(A-G) Body weight/composition, metabolic chambers, and GTT/ITT data n=7-20/group. Asterisks denote significant differences between diets: *p<0.05, **p<0.01, ***p<0.001, ****p<0.0001. (A-D) Šidák multiple comparison post 2-way ANOVA; (E-G) Dunnett multiple comparison post 2-way ANOVA

Figure S7

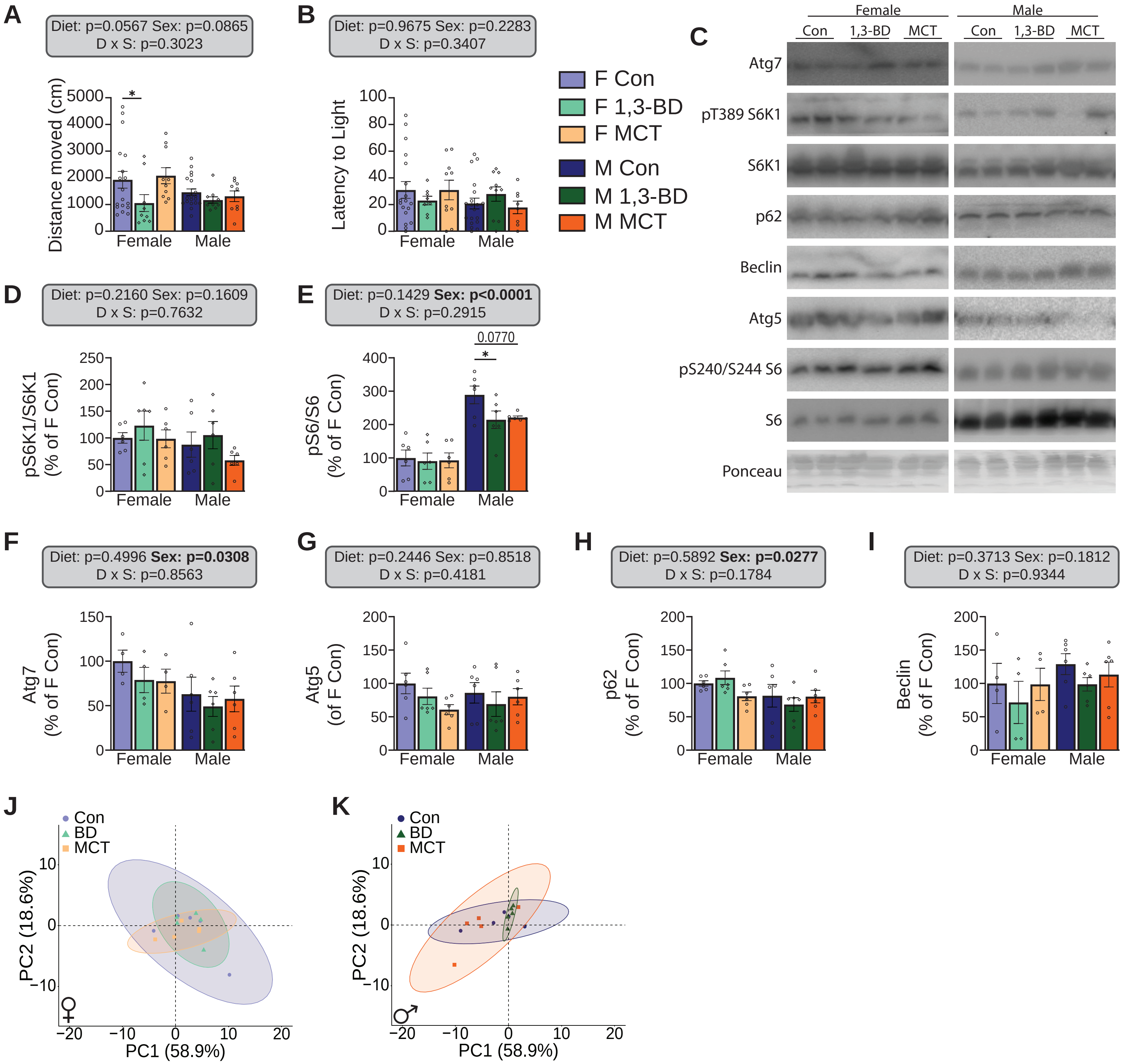

A
Diet: p=0.0567 Sex: p=0.0865
D x S: p=0.3023
Distance moved (cm)
Female
Male
B
Diet: p=0.9675 Sex: p=0.2283
D x S: p=0.3407
Latency to Light
F Con
F 1,3-BD
F MCT
M Con
M 1,3-BD
M MCT
C
Female
Male
Con
1,3-BD
MCT
Atg7
pT389 S6K1
S6K1
p62
Beclin
Atg5
pS240/S244 S6
S6
Ponceau
D
Diet: p=0.2160 Sex: p=0.1609
D x S: p=0.7632
pS6K1/S6K1 (% of F Con)
E
Diet: p=0.1429 **Sex: p<0.0001**
D x S: p=0.2915
0.0770
pS6/S6 (% of F Con)
F
Diet: p=0.4996 **Sex: p=0.0308**
D x S: p=0.8563
Atg7 (% of F Con)
G
Diet: p=0.2446 Sex: p=0.8518
D x S: p=0.4181
Atg5 (of F Con)
H
Diet: p=0.5892 **Sex: p=0.0277**
D x S: p=0.1784
p62 (% of F Con)
I
Diet: p=0.3713 Sex: p=0.1812
D x S: p=0.9344
Beclin (% of F Con)
J
Con
BD
MCT
PC2 (18.6%)
PC1 (58.9%)
K
Con
BD
MCT
PC2 (18.6%)
PC1 (58.9%)

**Figure S7: 1,3-BD and MCT reduce S6 phosphorylation.**

A) Distance moved in open field test.

B) Latency to light portion when placed in dark portion of dark/light box.

C-I) Immunoblotting for downstream targets of mTORC1 and autophagy markers (C). Quantification of pS6K/S6K (D), pS6/S6 (E), Atg7 (F; normalized to ponceau), Atg5 (G; normalized to ponceau), p62 (H; normalized to ponceau), and Beclin (I; normalized to ponceau).

J-K) PCA of brain polar metabolites in female (J) and male (K) mice.

(A-B) Behavioral assays n=7-20/group. (C-K) Histological and molecular assays n=4-6/group. Asterisks denote significant differences between diets: *p<0.05, **p<0.01, ***p<0.001, ****p<0.0001. (A-I) Dunnett multiple comparison post 2-way ANOVA

**Table S1:** Composition and calorie content for diets used in this study.

**Table S2:** Antibodies used for Western blotting and immunohistochemistry.
